# De novo design of protein nanoparticles with integrated functional motifs

**DOI:** 10.64898/2025.12.19.695620

**Authors:** Cyrus M. Haas, Sanela Rankovic, Hanul K. Lewis, Kenneth D. Carr, Connor Weidle, Sophie S. Gerdes, Lily R. Nuss, Felicitas Ruiz, Syed Moiz, Maggie Fiorelli, Emily Grey, Jackson McGowan, Nikhila Kumar, Adrian Creanga, Alex Kang, Hannah Nguyen, Yanqing Wang, Banumathi Sankaran, Annie Dosey, Rashmi Ravichandran, Asim K. Bera, Elizabeth M. Leaf, Cole A. DeForest, Masaru Kanekiyo, Andrew J. Borst, Neil P. King

## Abstract

Computational design of self-assembling proteins has long relied on pre-existing structures and sequences, fundamentally limiting control over their structural and functional properties. Recent machine learning-based methods have transformed our ability to design functional small de novo proteins and oligomers, yet methods to design large de novo protein assemblies with structures tailored to specific applications are still underexplored. Here, we develop a generalizable method for designing de novo symmetric protein complexes that incorporate target functional motifs into their structures. We report 34 new protein nanoparticles that form on-target assemblies with cubic point group symmetries. The nanoparticles exhibit a wide variety of backbones that were designed with atom-level accuracy, as evidenced by several cryo-EM and crystal structures that reveal minimal deviations from the design models. We use the method to generate a de novo antigen-tailored nanoparticle vaccine that elicits robust immune responses in mice. These results establish a generalizable approach that can be used to design functional self-assembling protein complexes with structures tailored to specific applications.

## Introduction

Large self-assembling protein complexes are ubiquitous in biology and perform a wide variety of specialized functions, from viral capsids protecting viral genomes to microtubules providing tracks for directional transport within cells (*1*, *2*). Their prevalence and sophistication has helped motivate the development of strategies to design novel protein assemblies for applications in medicine and beyond. Yet most approaches to date rely on using native structures found in the Protein Data Bank (PDB) (*3*) as building blocks for generating new self-assembling complexes (*4–7*). This fundamentally constrains our ability to design assemblies with user-defined structures, ultimately limiting their utility.

Considerable progress has been made in recent years towards developing de novo functional proteins. For example, machine learning-based tools like RoseTTAFold diffusion (RFdiffusion) (*8*), ProteinMPNN (*9*), and AlphaFold (*10*, *11*) have revolutionized the design of smaller proteins and protein complexes, including many de novo binders and small oligomers with functions ranging from antivirals (*12*) to receptor agonists (*13*) to targeted protein degradation (*14*). However, designing larger de novo self-assembling protein complexes with specific functions, like protein nanoparticles, has remained difficult. A small number of de novo protein nanoparticles have been generated using techniques such as reinforcement learning (*15*) or diffusion-based generative models (*8*). Others have been built by positioning pre-existing protein interfaces in three-dimensional (3D) space and filling the gaps between them with de novo protein backbone (*16*, *17*). Despite these advances, no methods have yet been reported for protein nanoparticle design that enable full control over the position of every atom, allow the incorporation of user-defined functional motifs, and generalize to any symmetry group.

Here, we establish a computational method to design de novo self-assembling protein nanomaterials bearing functional motifs that are tailored to specific applications. We demonstrate how this method can be applied to generate nanoparticle scaffolds customized for display of an influenza virus glycoprotein antigen, and find that the resultant nanoparticle vaccine candidate elicits robust antibody responses in mice.

## Results

### Computational design of de novo nanoparticles

We selected homomeric (i.e., “one-component”) nanoparticles with tetrahedral, octahedral, and icosahedral point group symmetry as demanding targets of our ML-based approach to de novo self-assembling protein nanomaterials design. Most recent successes in novel protein nanoparticle design have been achieved using some variation of the following strategy: identify naturally occurring or previously designed oligomeric protein building blocks, dock those oligomers into a target symmetric architecture, and design a new protein-protein interface between the docked building blocks to drive nanoparticle assembly (*4*, *5*, *7*, *18*, *19*). We posited that the dependency on pre-existing structures could be eliminated by leveraging newer protein design approaches to build nanoparticles from de novo protein components instead, unlocking a conceptually limitless design space (**Fig. 1**). We selected as our first target one-component icosahedral nanoparticles consisting of twenty trimeric building blocks (the “I3” architecture). We used RFdiffusion to generate a library of approximately 30,000 de novo trimers, with each protomer consisting of either 150 or 200 amino acids (**Fig. 1A**). These components were then docked according to icosahedral symmetry using RPXdock (*20*) followed by additional sampling of the translational degree of freedom using the Rosetta SymDofMover (**Fig. 1B**, (*18*)). The sequence of the entire nanoparticle, comprising all interactions within and between the 60 protomers, was then designed in one step with ProteinMPNN (**Fig. 1C**). Finally, we predicted the structures of the designed oligomers with AlphaFold2 (AF2) to identify those in which the designed amino acid sequence strongly encoded the target structure (**Fig. 1D**). Metrics from both AF2 (pLDDT > 90, pAE < 5) and Rosetta-based evaluation of nanoparticle design models (ddG < −10) were used to filter the designs, and passing designs were further down-selected by visual inspection. Encouraged by the high quality of the designs in silico, we also generated tetrahedral nanoparticles built from six de novo dimers (“T2”) and octahedral nanoparticles built from six de novo tetramers (“O4”) (see *Materials and Methods*). We selected 166 icosahedral, 84 tetrahedral, and 56 octahedral nanoparticles to evaluate experimentally.

**Figure 1.**
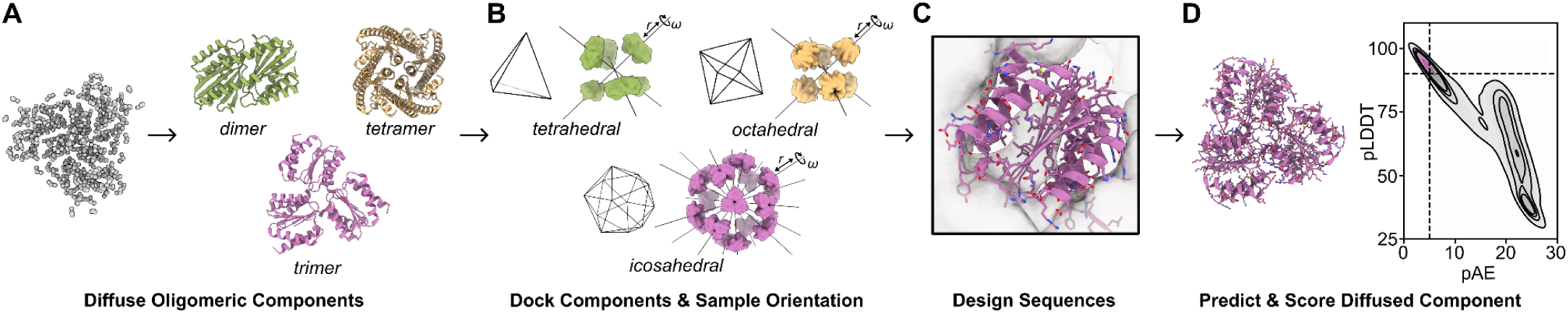
Overview of the design pipeline for generating de novo protein nanoparticles. (**A**) RFdiffusion is used to generate libraries of cyclic oligomers that are used as inputs for RPXdock. (**B**) Rotational (*ω*) and translational (*r*) degrees of freedom are sampled by RPXdock to identify optimal positioning of components in tetrahedral, octahedral, or icosahedral symmetry. (**C**) ProteinMPNN is used to design the sequence of the entire homomeric nanoparticle in one step. (**D**) AF2 structure prediction of the designed oligomers is used to identify constructs for experimental characterization. Dashed lines on the plot represent pLDDT = 90 and pAE = 5 and designs in the upper left quadrant (pink) are advanced to ddG scoring and visual inspection.

### Screening and characterization of de novo protein nanoparticles

Genes encoding the one-component nanoparticles with appended hexahistidine tags were synthesized for expression in *Escherichia coli* and the resultant proteins purified from clarified lysates by immobilized metal affinity chromatography (IMAC). After small-scale expression and high-performance liquid chromatography (HPLC), putative hits were identified and scaled up for detailed characterization. Of the tested constructs, we identified a total of 2 tetrahedral nanoparticles (**Fig. 2A**), 4 octahedral nanoparticles (**Fig. 2B**, and fig. S1, A and B), and 16 icosahedral nanoparticles (**Fig. 2C**, and fig. S1, C to P) that formed the expected on-target assembly. Each of the tetrahedral and octahedral nanoparticle subunits featured unique folds, and 13 of the 16 icosahedral nanoparticles had distinct backbones, though all sequences were unique. These 19 distinct protein backbones comprised a diverse set of secondary and tertiary structures, reflecting the ability of generative methods like RFdiffusion to create highly diverse structures. Preparative size exclusion chromatography (SEC) of the IMAC-purified nanoparticles revealed a range of assembly behaviors: some were highly monodisperse (**Fig 2C**, and fig. S1, C and D), others exhibited shoulders consistent with aggregation or alternative assembly states (**Fig. 2A**, and fig. S1, B, E to J), and several showed clear signs of aggregation, unassembled component, or both (**Fig. 2B**, and fig. S1, A, K to P). Most of the SEC-purified nanoparticles appeared monodisperse by dynamic light scattering (DLS), with a few exceptions that indicated some aggregation (**Fig. 2** and fig. S1). Finally, we used negative stain electron microscopy (nsEM) to visualize the nanoparticles and found that most samples were homogeneous, though after extensive nsEM classification and processing, some off-target species were also observed in 31% of icosahedral samples (fig. S2). The on-target densities obtained by 3D reconstruction closely match the design models in every case (fig. S2). Finally, for the tetrahedral and octahedral nanoparticles, some disassembly was observed in nsEM micrographs (**Fig. 2A** and fig. S1A).

**Figure 2.**
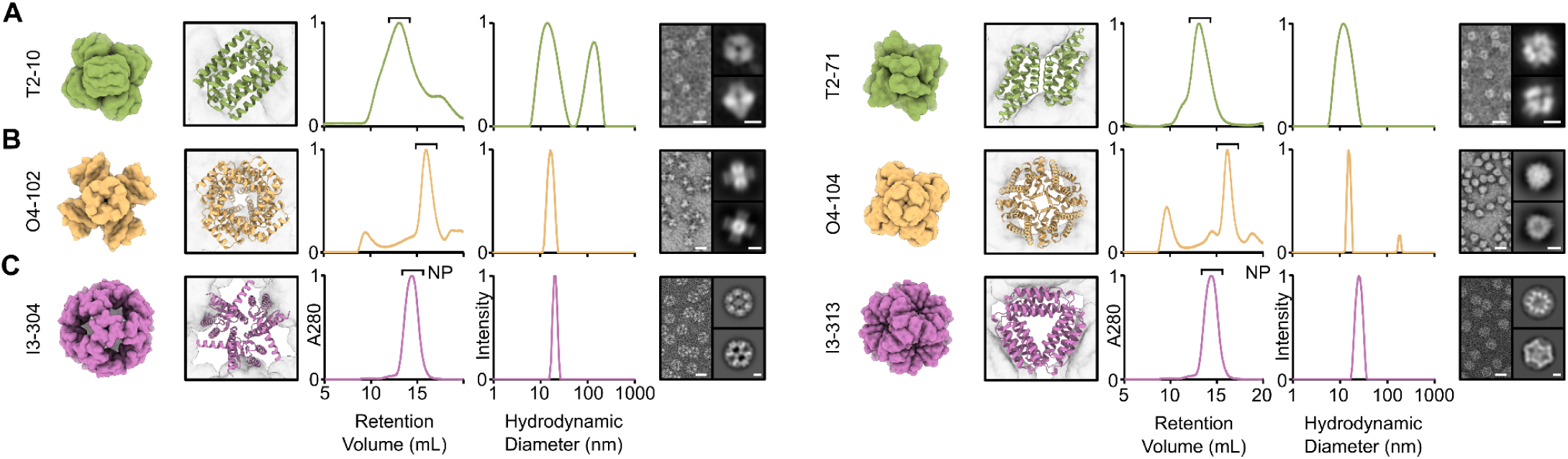
Experimental characterization of de novo nanoparticles. From left to right, design models, preparative SEC, DLS, nsEM micrographs, and 2D class averages are shown for (**A**) two tetrahedral nanoparticles (T2-10 and T2-71), (**B**) two octahedral nanoparticles (O4-102 and O4-104), and (**C**) two icosahedral nanoparticles (I3-304 and I3-313). Volume maps of the nanoparticle design models are shown with an enlarged crop of the diffused oligomer (dimer, tetramer, and trimer) to the right of each model. SEC and DLS plots are normalized and nanoparticle fractions purified for further characterization are indicated (“NP”). Scale bars are 20 nm for the micrographs and 5 nm for the 2D class averages.

### High-resolution structure determination of de novo protein nanoparticles

We selected one icosahedral (I3-304) and two octahedral (O4-102 and O4-104) nanoparticles for cryo-electron microscopy (cryo-EM) structure determination to evaluate the accuracy of our method at high resolution. Micrographs for all three materials contained clearly defined particles, and 2D class averages revealed distinct views for each design consistent with their respective icosahedral or octahedral geometries that captured all major symmetry axes and additional orientations for each (**Fig. 3A**). We obtained three-dimensional (3D) reconstructions at global resolutions of 2.14 Å, 2.14 Å, and 2.40 Å for I3-304, O4-102, and O4-104, respectively (**Fig. 3A**, fig. S3-6 and Table S2). For I3-304, we also observed and characterized a minor off-target species found in 1% of selected particles (fig. S4 and Table S3). The alpha-carbon (Cɑ) root mean square deviations (RMSD) between all 60 or 24 subunits of the design models and the cryo-EM structures were 1.7 Å, 1.0 Å, and 1.7 Å, respectively (**Fig. 3B**). Cɑ RMSDs for two trimeric or tetrameric oligomers across each two-fold interface were 1.3 Å, 1.0 Å, and 1.5 Å (**Fig. 3C**), while comparing individual trimers or tetramers resulted in Cɑ RMSDs of 1.2 Å, 0.9 Å, and 0.9 Å (**Fig. 3D**) for I3-304, O4-102, and O4-104, respectively. We also determined a crystal structure of one tetrahedral nanoparticle, T2-71, at a resolution of 3.2 Å (Table S4). The refined structure matches the design model closely, with Cɑ RMSDs of 1.7 Å for the full nanoparticle, 1.0 Å for three subunits surrounding the three-fold interface, and 1.4 Å for the two subunits forming the dimer (**Fig. 3E**). Qualitative analysis of the interfaces of the designed assemblies revealed that the side chains at the two-fold (nanoparticle) interface of I3-304 resembled the hydrophobic packing frequently observed in previous nanoparticle design approaches (**Fig. 3F**, (*7*, *18*, *19*, *21*)). However, analysis of O4-102 and O4-104 revealed that the four-fold interfaces within each tetrameric building blocks are stabilized by a core of hydrophobic residues surrounded by peripheral polar interactions, while the two-fold interfaces between the building blocks depend largely on low-resolution backbone shape complementarity with more limited and more polar side-chain interactions (**Fig. 3G**). In T2-71, both the two-fold and three-fold interfaces have an approximately even balance of hydrophobic and polar interactions (**Fig. 3H**). These observations motivated analysis of the design models for all designs selected for experimental characterization. Assuming the close agreement we observed between all four designed and experimentally determined structures holds, the oligomeric interfaces within the diffused building blocks comprised on average slightly more than 50% hydrophobic residues across all architectures (fig. S7). However, the nanoparticle assembly interfaces exhibited more variation in their hydrophobicity, with the icosahedral designs favoring more hydrophobic interfaces, the octahedral designs skewing less hydrophobic, and the tetrahedral designs yielding a bimodal distribution (fig. S7).

**Figure 3.**
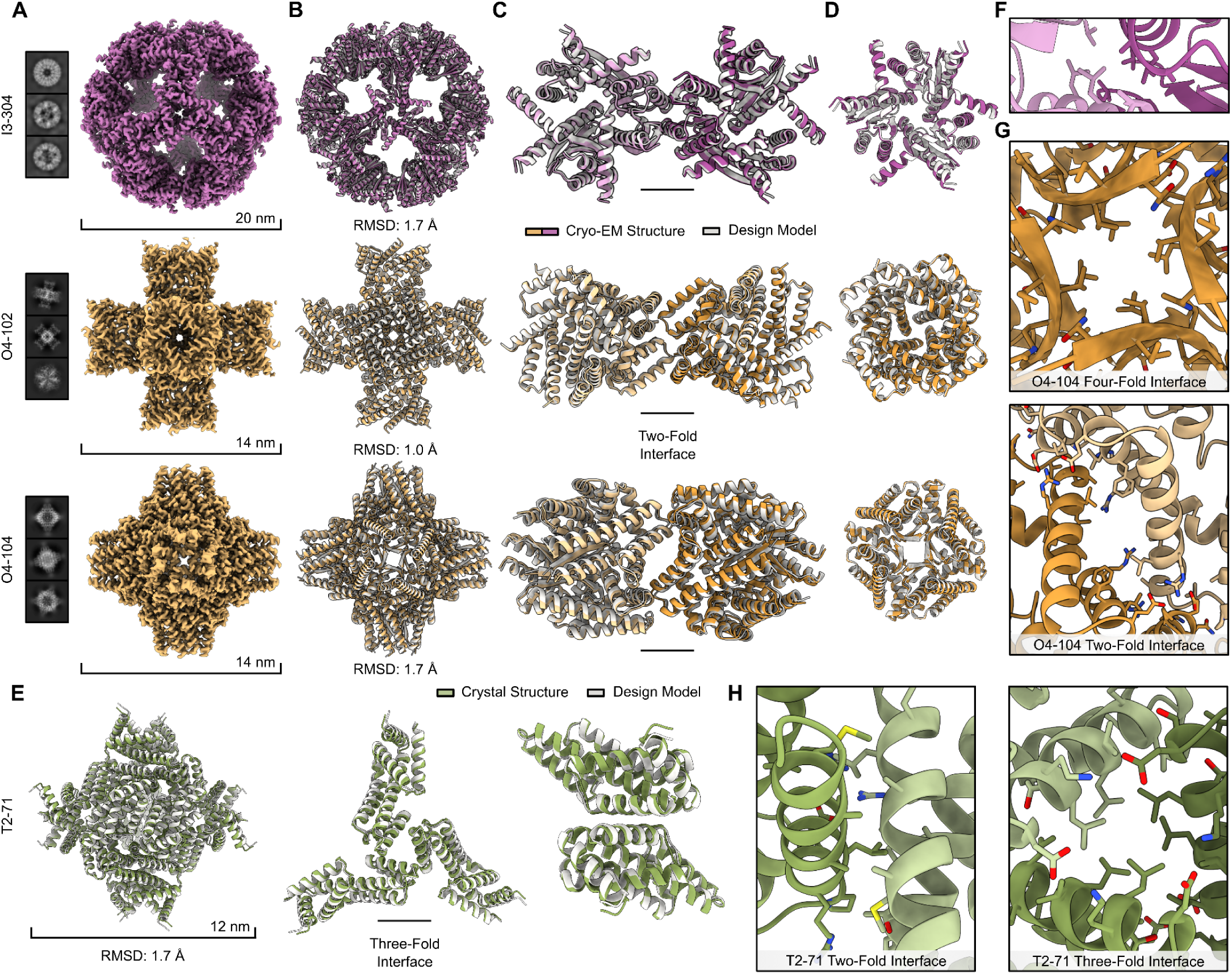
High-resolution structure determination of four de novo protein nanoparticles. (**A**) Cryo-EM 2D class averages and 3D reconstruction volume maps of three protein nanoparticles. (**B**) Overlays of and Cɑ RMSDs between the design models (gray) and cryo-EM structures (purple or yellow). (**C**) Overlays and RMSDs of the design models and cryo-EM structures for two oligomeric building blocks surrounding the two-fold symmetry axis. (**D**) The designed trimeric or tetrameric oligomer for each nanoparticle overlaid with the corresponding cryo-EM structure. (**E**) Overlay of the design model (gray) and crystal structure (green) for the tetrahedral nanoparticle T2-71, its primary the three-fold interface, and the dimeric component. (**F**) Details of the two-fold interface in the cryo-EM structure of I3-304. (**G**) (top) Details of the four-fold tetrameric interface and (bottom) two-fold nanoparticle interface in the cryo-EM structure of O4-104. (**H**) (left) Details of the two-fold dimeric interface and (right) the primary three-fold nanoparticle interface in the crystal structure of T2-71.

### Designing tailored protein nanoparticles for specific applications

Encouraged by the success of the design pipeline described above, we wanted to test whether it could be extended to design protein nanoparticles with structures tailored to a specific application. We used the de novo design of antigen-tailored nanoparticle immunogens as a demanding test case. Multivalent display of antigens on the surface of nanoparticle scaffolds is a widely used approach to improve the magnitude, quality, and duration of vaccine-elicited immune responses (*22–27*), as repetitive antigen display enhances B-cell receptor clustering and B cell activation (*28–30*), while nanoparticle size improves trafficking of the immunogen in vivo (*31*). Three recently licensed vaccines for malaria (*32*, *33*) and COVID-19 (*34*) have employed this approach, yet the fixed structures of protein nanoparticle scaffolds used to date ultimately limits its generalizability. For example, there are several classes of antigens that are promising vaccine targets but are difficult to display on nanoparticle scaffolds due to symmetry mismatches or geometric constraints. These include, among others, tetrameric antigens with C4 symmetry, heterodimeric antigens, and antigens that already form larger assemblies. To test whether we could custom-build de novo nanoparticle scaffolds for specific antigens, we selected the influenza virus hemagglutinin (HA) “trihead” antigen: a computationally designed, closed receptor binding domain trimer, stabilized with a disulfide bond and fused to the trimeric coiled-coil GCN4 (*35*).

To design trihead-tailored nanoparticles, we used symmetric motif scaffolding in RFdiffusion to generate approximately 200,000 trimeric building blocks containing the C-terminal seven residues from the base of the coiled-coil domain of the antigen (**Fig. 4A**). The rest of the pipeline stayed essentially the same as before: we docked the de novo trimers into icosahedral symmetry using RPXdock; designed sequences for the entire nanoparticles in one step with ProteinMPNN; and predicted the structures of the designed trimers with AF2. The sequence of the coiled-coil motif was allowed to vary during design to increase the likelihood of its successful incorporation into the de novo component. After visually inspecting designs that passed AF2 and Rosetta metrics (pLDDT > 90, pAE < 5, ddG < −10), we selected 180 designs to test experimentally. Genes encoding the designed nanoparticles were synthesized, first without antigen to enable easier screening in bacterial cells. After small-scale expression in *E. coli* followed by HPLC of IMAC-purified protein, we selected 16 putative hits for scaled-up expression and characterization, which revealed that 12 designs (from 11 unique backbones) formed expected icosahedral assemblies (**Fig. 4B** and fig. S8). Like before, the backbones of these designs were quite diverse, though they all contained the target antigen-derived motif at the icosahedral three-fold axis of symmetry (**Fig. 4C**). All of the nanoparticles yielded preparative SEC traces indicating some aggregation and some, like I3-320, have clear unassembled component peaks as well (**Fig. 4D** and fig. S8). Nevertheless, DLS of the SEC-purified nanoparticle fractions generally suggested monodispersity with the exception of I3-326, I3-330, and I3-331, which all contained small amounts of aggregates (**Fig. 4E**, and fig. S8). nsEM of the nanoparticles (**Fig. 4F**, and fig. S8) confirmed that most were homogenous and formed the intended structures, though some off-target assemblies were also observed (fig. S9).

**Figure 4.**
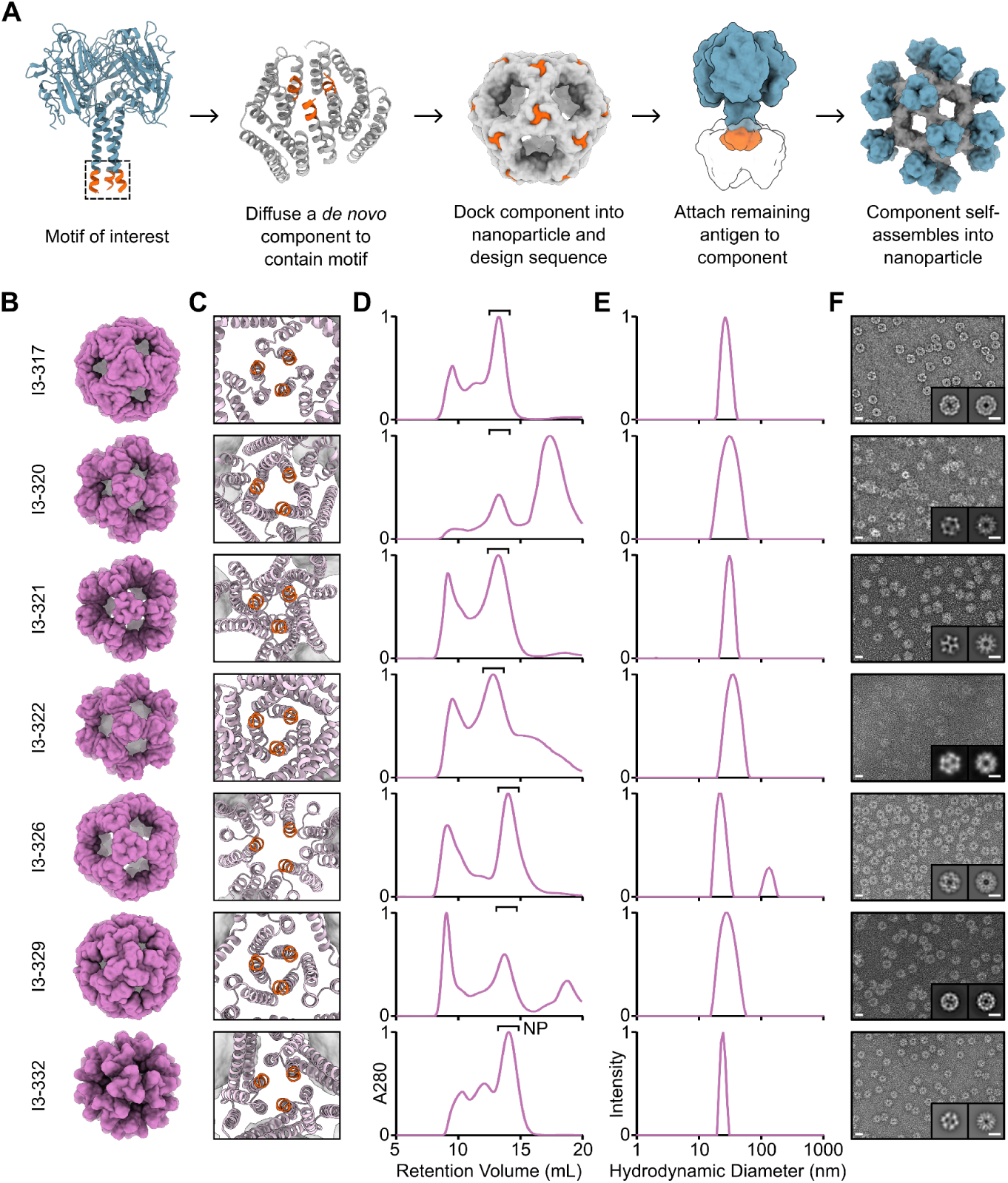
Antigen-tailored nanoparticle design and characterization. (**A**) An overview of the design pipeline for building antigen-tailored one-component protein nanoparticles. (**B**) Design models for a subset of successfully assembling tailored protein nanoparticles. (**C**) A close-up of the trimeric interface of each protein nanoparticle, highlighting the scaffolded motif in orange. (**D**) Preparative SEC traces for each nanoparticle. Fractions purified for further characterization are indicated (“NP”). (**E**) DLS of the SEC-purified nanoparticles. (**F**) nsEM micrograph and 2D-class averages for each nanoparticle. Scale bars = 20 nm and 10 nm for micrographs and 2D-class averages, respectively.

### Antigen display and immunogenicity

Having demonstrated successful assembly of the antigen-tailored nanoparticles in *E. coli*, we next tested if the nanoparticles could secrete from mammalian cells with the trihead antigen displayed. We ordered genes encoding each nanoparticle with the HA trihead from an H1N1 influenza virus genetically fused to the N terminus. We identified one nanoparticle that secreted (based on I3-326), and scaled up expression of that nanoparticle displaying the trihead of the recent H1N1 vaccine strain A/Michigan/45/2015 (MI15) for detailed characterization. I3-326-MI15-TH secreted as an assembled nanoparticle immunogen, yielding SEC, DLS, and nsEM data that revealed a high degree of monodispersity (**Fig. 5A**). Additionally, antigenic characterization via biolayer interferometry (BLI) showed binding of the monoclonal antibody 5J8 to the receptor binding site and no binding of the trimer interface-directed antibody FluA-20, indicating display of closed antigen trimers as intended (**Fig. 5B**). We further structurally characterized I3-326-MI15-TH using cryo-EM. Electron micrographs revealed homogenous, monodisperse particles, and 2D classification of the immunogen showed clearly defined HA trihead antigens repetitively displayed on the I3-326 surface (**Fig. 5C**). We obtained a 3D reconstruction at a global resolution of 3.1 Å (**Fig. 5D** and fig. S10). When compared to the design model, the cryo-EM structure of the nanoparticle had a Cɑ RMSD of 3.0 Å (**Fig. 5E**), which is slightly larger than the deviations observed for other nanoparticles described here. The scaffold exhibits a subtle overall bowing (**Fig. 5F**), likely a result of its entirely alpha-helical architecture and hinging at the loop junctions. Even so, the RMSD between the design model and cryo-EM structure of the trimeric building block is still relatively low at 1.9 Å, and the antigen-derived motif was accurately scaffolded at the three-fold axis of the building block. The lower resolution of the displayed antigen indicated some flexibility in the linker connecting it to the nanoparticle.

**Figure 5.**
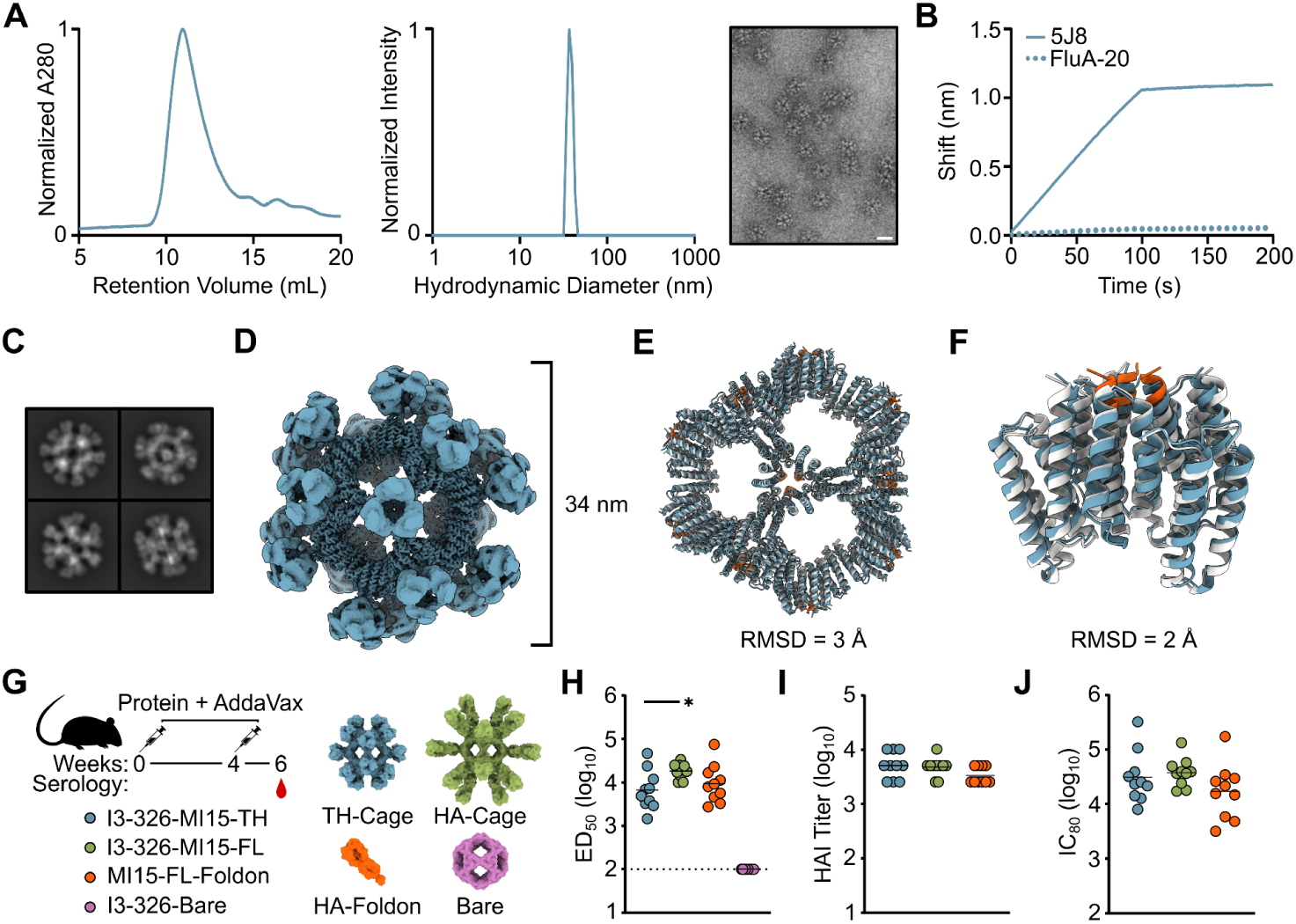
Immunogen characterization and immunogenicity. (**A**) SEC, DLS, and nsEM for the lead nanoparticle immunogen I3-326-MI15-TH. Scale bar = 34 nm. (**B**) BLI for nanoparticle I3-326-MI15-TH. (**C**) Cryo-EM 2D class averages of I3-326-MI15-TH. (**D**) 3D reconstruction of I3-326-MI15-TH. (**E**) Overlay of the cryo-EM structure of the nanoparticle (blue) with the design model (gray). The scaffolded motif is shown in orange on the design model. (**F**) Overlay of one trimer from the cryo-EM structure with the design model, colored as in (**G**) Immunogenicity study design and immunogens tested. (**H**) Week 6 ELISA ED_50_ titers, (**I**) HAI titers, and (**J**) IC_80_ neutralization titers. All statistical significance is shown (not including negative control) and was determined using the Kruskal-Wallis test corrected with Dunn’s multiple comparisons test.

We next tested I3-326-MI15-TH as a protein nanoparticle immunogen in vivo. We also evaluated the same nanoparticle scaffold displaying a full-length, trimeric HA ectodomain. This nanoparticle, I3-326-MI15-FL, was well behaved, monodisperse within the selected fraction, and antigenically intact (fig. S11). We also included the full-length trimeric HA ectodomain genetically fused to the T4 foldon trimerization domain and the bare I3-326 nanoparticle, I3-326-Bare, as positive and negative controls, respectively (**Fig. 5G**). Groups of 10 BALB/c mice were each injected with molar equivalents of 1 µg of HA trihead antigen, all adjuvanted with AddaVax. They were boosted at week four and serum was collected at week six. HA-specific IgG titers were measured by ELISA against full-length HA ectodomain trimer on a different trimerization domain, I53_dn5B. All positive groups yielded comparable titers, with the I3-326-MI15-FL nanoparticle eliciting slightly higher titers when compared to the trihead nanoparticle, as expected (**Fig 5H** and fig. S12). When examining the functional activity of vaccine-elicited antibodies with a hemagglutination inhibition (HAI) assay, no statistically significant differences were observed between groups (**Fig 5I**). A similar trend was observed for vaccine-matched neutralization, where all immunogens exhibited similar IC_80_ values (**Fig 5J**). These data demonstrate that the de novo nanoparticle pipeline used to design antigen-tailored nanoparticles can successfully generate immunogens that elicit robust antibody responses in vivo.

## Discussion

Here we describe a robust computational strategy for the de novo design of functional, self-assembling protein nanomaterials. The central methodological advance is the use of RFdiffusion to generate de novo oligomeric protein building blocks that are docked into target architectures prior to amino acid sequence design. This approach enables precise atom-level specification of the entire assembly and removes the dependence on pre-existing building blocks or interfaces, a constraint that has limited customization in many previous methods. Successful application to three geometrically demanding architectures (T2, O4, and I3) demonstrates that, like earlier strategies, the approach generalizes across point-group symmetries (*4–7*, *18*, *19*, *21*). Notably, despite the increased difficulty of designing assemblies entirely de novo, we observed success rates comparable to approaches that rely on pre-existing components (>10% for icosahedral assemblies).

Beyond these advances, the method differs from prior approaches in two important respects. First, the resulting assemblies comprise subunits spanning a wide range of secondary and tertiary structures, consistent with RFdiffusion’s generative capacity and suggesting broad potential for scaffolding diverse structural and functional motifs. Second, the approach is rapidly deployable given sufficient computational resources: several independent design campaigns produced experimentally validated de novo assemblies within eight weeks of initiating design.

To date, most applications of protein nanoparticles have relied on modifying pre-existing assemblies—either natural or computationally designed—to confer new functions. In vaccine design, for example, antigens are typically “retrofitted” onto established nanoparticle scaffolds that may or may not provide an optimal geometric match (*22–25*). The approach described here inverts this paradigm by using a functional motif as the seed from which custom nanoparticle subunits are grown. This strategy is enabled by the robustness of motif scaffolding in RFdiffusion and related methods, which have previously been used to generate de novo monomers and small oligomers containing defined functional elements (*12–14*, *36*, *37*). Here we extend motif scaffolding for the first time to the design of de novo protein nanoparticles tailored to a specified function.

Although our initial demonstration focused on nanoparticles whose building blocks are geometrically optimized for genetic fusion to a specific antigen, the same conceptual framework should enable precise control over a wide range of nanomaterial properties, including size, shape, symmetry, and the placement of sites for binding, catalysis, post-translational modification, and other functions. Given the demonstrated versatility of RFdiffusion in designing functional monomers and oligomers, we anticipate that this approach will enable comparably versatile design of functional de novo protein assemblies.

The coiled-coil motif we scaffolded to generate the I3-326-MI15-TH nanoparticle immunogen is shared by multiple families of viral glycoprotein antigens, including the fusion (F) proteins of paramyxoviruses such as Hendra, measles, and mumps viruses, suggesting I3-326 may provide a generalizable scaffold for multivalent display of antigens from these families (*38*, *39*). We note that among the 12 motif scaffolded de novo nanoparticles evaluated, we only detected secretion of I3-326 fused to the MI15-TH glycoprotein antigen during small-scale screening. This observation is consistent with recent reports indicating that secretion of protein nanoparticles and nanoparticle immunogens is a poorly understood and idiosyncratic phenomenon (*40*, *41*). Nevertheless, I3-326-MI15-TH was secreted as a monodisperse, antigenically intact nanoparticle and elicited robust immune responses in mice.

Although we designed only one-component functional assemblies here, the generality of the approach suggests that it should extend readily to more sophisticated, multi-component protein nanomaterials. A remaining limitation lies in the generation of suitable de novo backbones using RFdiffusion, which currently requires sampling tens to hundreds of thousands of candidate backbones to identify building blocks that dock together well in a target architecture. We expect that future advances in machine-learning methods for protein design will address this by enabling optimization of the shape complementarity of subunits during generation of entire assemblies. Also important will be the development of improved computational filters and evaluation metrics that more directly predict successful protein nanoparticle assembly. The assemblies and methodology reported here provide an opportunity to generate large, systematic data sets that can be used to define such metrics.

## Data, Materials, and Software Availability

Data deposition, atomic coordinates, and structure factors have been deposited in the Protein Data Bank, http://www.rcsb.org (PDB ID 9ZQI, 9ZOJ, 9ZOL, 9ZPM, 9ZSD, 9ZV3). Cryo-EM maps were deposited in the Electron Microscopy Data Bank (EMD-74566, EMD-74498, EMD-74499, EMD-74530, EMD-74706). These data will be released prior to publication. Design models for all ordered and experimentally validated nanoparticle constructs can be downloaded using the following stable link: https://files.ipd.uw.edu/pub/cyhaas/haas_denovo_nanoparticles.zip.

## Acknowledgements

The authors thank Suna Cheng and Adri Tran-Pearson for assistance with tissue culture; Kandise VanWormer, Hernan Nunez-Ortega, and Eryn Weston for laboratory and administrative support; Luki Goldschmidt, Patrick Vecchiato, and Bulat Faezov for building and maintaining the computing infrastructure at the Institute for Protein Design; and Joel Quispe, Fangfang Zhang, and Wentao Jiang for management of the University of Washington Cryo-EM facilities. This work was supported by the National Institute of Allergy and Infectious Disease (1P01AI167966, U19AI181881, and U54AI170856 to N.P.K.); the Advanced Research Projects Agency for Health (ARPA-H; APECx Program Award No. 1AY1AX000036-01 to N.P.K.); the Gates Foundation (INV-069938 to N.P.K.); a generous gift from Coefficient Giving; and the National Science Foundation Graduate Research Fellowships Program Award (DGE-2140004 to C.M.H.). This study was supported in part by the Intramural Research Program of the National Institutes of Health (NIH) (M.K.). The contributions of the NIH authors are considered works of the U.S. Government. The findings and conclusions presented in this paper are those of the authors and do not necessarily reflect the views of the NIH or the U.S. Department of Health and Human Services. This research used resources (FMX/AMX) of the National Synchrotron Light Source II, a US Department of Energy (DOE) Office of Science User Facility operated for the DOE Office of Science by Brookhaven National Laboratory under contract DE-SC0012704. The Center for BioMolecular Structure (CBMS) is primarily supported by the National Institutes of Health, National Institute of General Medical Sciences (NIGMS) through a Center Core P30 grant (P30GM133893), and by the DOE Office of Biological and Environmental Research (KP1607011). Cryo-EM facilities and resources were provided by the University of Washington Arnold and Mabel Beckman cryo-EM center and the cryo-EM facility at HHMI Janelia Research Campus.

## Competing Interests Statement

C.M.H., S.R., A.D., and N.P.K. are coinventors on a patent application related to this work. All other authors declare no competing interests.

## Materials and Methods

### Designing de novo one-component protein nanoparticles

RFdiffusion was used to generate libraries of de novo dimers, tetramers, and trimers to create tetrahedral (T2), octahedral (O4), and icosahedral (I3) protein nanoparticles, respectively. All protomers were either 150 or 200 amino acids in length–both lengths were tested for all architectures. For motif-scaffolded icosahedral nanoparticles, RFdiffusion was also used but with the desired motif as an input. With these oligomers as input, we ran RPXdock with the standard score function (for T2, O4, and I3) and the SASA score function (for T2 and O4). We then filtered docks using the RPX score (standard > 60 for T2 and O4, standard > 70 for I3, sasa > 5 for O4, sasa > 6 for T2) to identify more optimal docks. At this stage, we used the Rosetta SymDofMover to further sample the single translational degree of freedom and expand the available set of nanoparticles for sequence design. We then filtered for clashes, considered any Cɑ contacts closer than 3 Å (for O4 and I3) or 2.5 Å (for T2), and number of contacts, considered as Cɑ contacts between 3-8 Å (≥10 contacts required for all architectures). The docks that passed these filters were saved as whole nanoparticle PDB files and sent through ProteinMPNN, where we designed 2-8 sequences for each nanoparticle. The sequences were filtered to remove any that contained alanine parties (defined as four or more consecutive alanines) and to remove any sequences that were predicted to contain a cryptic transmembrane region using the Rosetta degreaser metric (ddG_ins,pred_ < 2.6) (*40*). The remaining sequences were predicted with AlphaFold2 multimer_v3 in the original oligomeric state in which they were diffused. We downselected constructs yielding pLDDT > 90 and pAE < 5 and threaded the passing sequences back onto their parent nanoparticle docks. Rosetta scripting was used to generate the design model from the threaded nanoparticle dock asymmetric unit and calculate ddG. We used a ddG cutoff of −10, followed by visual inspection of all remaining outputs to select designs for experimental characterization.

### *E. coli* expression and purification

Synthetic genes encoding the designs procured from IDT or Twist and assembled via Golden Gate Assembly were transformed into BL21 competent *E. coli* cells (NEB). For screening constructs at 1 mL scale, cells were grown in 96-deep well format in autoinduction media (Terrific Broth II media supplemented with 50×5052, 50×M salts, 20 mM MgSO_4_, and a trace metal mix) at 37°C for 24 hours at 1350 revolutions per minute (rpm) in a Heidolph shaker. Large-scale 50 mL and 500 mL cultures were grown in Terrific Broth II media (MP Bio) and induced with 0.1 mM IPTG at OD_600_=0.6-0.8 at 37°C and grown for 12-15 hours at 37°C and 250 rpm. Cells were then spun down at 4000 *g* for 30 minutes and resuspended in lysis buffer (25 mM Tris pH 8, 250 mM NaCl, 20 mM Imidazole, 200 mM L-Arginine) with the addition of bovine pancreatic DNase I (Sigma-Aldrich) and protease inhibitors (Thermo Scientific). Resuspended cells were then sonicated with either a 24-prong sonicator (for 96-well format) or a 4-prong sonicator (for larger expressions). Lysed cells were centrifuged for 30 minutes at 4000 *g* and clarified lysate was subsequently poured over equilibrated Ni-NTA resin (QIAGEN) in a gravity column. The columns were washed with 10 column volumes of wash buffer (25 mM Tris pH 8, 250 mM NaCl, 20 mM Imidazole, 200 mM L-Arginine), then mixed with and incubated in elution buffer (25 mM Tris pH 8, 300 mM NaCl, 500 mM Imidazole, 200 mM L-Arginine) for 10 minutes prior to being eluted. Eluant was concentrated to 1 mL and further purified using Superdex 200 10/300 (for T2 architecture) or Superdex 6 10/300 (for O4 and I3 architectures) columns on an ÄKTA Pure (Cytiva) and buffer exchanged into a standard sizing buffer (25 mM Tris pH 8, 500 mM NaCl, 100 mM L-Arginine). Instrument control and elution profile analysis were performed using UNICORN 7 software (Cytiva).

### HEK293F expression and purification

Plasmids for mammalian expression were purified from DH5α competent *E. coli* (NEB) according to the QIAGEN Plasmid Plus Maxi Kit protocol (QIAGEN). For mammalian expression of secreted protein nanoparticles, Expi293F were transfected at 3 million cells/mL with the following mixture per mL of cells: 70 µL Opti-MEM, 1 µg plasmid DNA, and 3 µL polyethylenimine (PEI). After growth at 37°C, 8% CO_2_, 150 rpm for 3 days, the cells were centrifuged at 4000 *g* for 10 minutes before collecting and filtering the supernatant with a 0.2 µm filter. Equilibrated Ni-Sepharose Excel resin (Millipore Sigma) was then added to the supernatant and agitated for one hour at room temperature. The resin was then collected and washed with 20 column volumes of wash buffer (50 mM Tris, 500 mM NaCl, 30 mM Imidazole). The resin was then incubated in 5 column volumes of elution buffer (50 mM Tris, 500 mM NaCl, 300 mM Imidazole) for 10 minutes prior to elution. The elution step was repeated prior to concentrating the eluant to 1 mL and further purification was performed with a Superdex 6 10/300 column on an ÄKTA Pure (Cytiva) using 25 mM Tris pH 8, 500 mM NaCl, 100 mM L-Arginine as the running buffer.

### Dynamic light scattering

Proteins were analyzed on an Unchained Labs Uncle machine by adding 8.8 µL of SEC-purified protein samples at approximately 0.1 mg/mL into a quartz capillary cassette (Uni, Unchained Laboratories) and running a standard sizing and polydispersity protocol at 25°C. Buffer conditions were appropriately accounted for in the Uncle software.

### Negative stain electron microscopy

Protein samples were diluted to 0.10 mg/mL in 25 mM Tris pH 8.0, 500 mM NaCl, 100 mM L-arginine, and 5% (v/v) glycerol, except for I3-315 and I3-317, which were prepared at 0.033 mg/mL and 0.05 mg/mL, respectively. Aliquots (3 µL) were applied to glow-discharged 300-mesh carbon-coated copper grids (Electron Microscopy Sciences, CF-300-Cu-50; 30 s at 15 mA) and allowed to adsorb for 30 s before excess liquid was removed by wicking with filter paper. Grids were immediately stained with 2% (w/v) uranyl formate, with stain application and blotting repeated three times, followed by air drying after the final blot. Negative-stain data were collected on a 120 kV Talos L120C transmission electron microscope (Thermo Fisher Scientific) equipped with a BM-Ceta camera using a 1 s exposure time. Images were recorded at 57,000× magnification, corresponding to a pixel size of 2.49 Å, with defocus values ranging from −1.3 µm to −2.3 µm sampled in six steps. Data collection was automated using E Pluribus Unum (EPU; Thermo Fisher Scientific). Image processing was performed in cryoSPARC (v4.7.0 and v4.7.1; Structura Biotechnology Inc.). Micrographs were imported and subjected to patch-based CTF estimation. For particle picking, an initial subset of micrographs (approximately 20%) was used to seed blob-based particle picking with particle diameters ranging from 140 to 400 Å. In cases where blob picking yielded insufficient particles, manual picking was performed. Extracted particles (box sizes 250–300 pixels) were aligned and subjected to 2D classification, and well-resolved class averages corresponding to both on-target and off-target assemblies were selected. These class averages were used to generate ab initio 3D reconstructions with a maximum target resolution of 20 Å and appropriate symmetry applied, followed by homogeneous refinement to obtain final negative-stain volumes. Design models were docked into the corresponding density maps using UCSF ChimeraX to assess agreement between the computational designs and experimentally determined reconstructions. Off-target assemblies were processed using the same workflow as on-target particles, with additional validation steps. All off-target species were confirmed as present over ≥2 technical replicates. Ambiguous or poorly defined 2D class averages were omitted from all analyses to ensure the generation of reliable 3D volumes of off-target species. For symmetry determination, ab initio reconstructions of off-target particles were first generated without imposed symmetry and visually inspected to determine the appropriate symmetry. This was followed by homogeneous refinement with the identified symmetry operators enforced. Template projections generated from the refined off-target volumes were cross-validated against experimental 2D class averages to assess reconstruction accuracy and angular coverage. Off-target volumes exhibiting trimeric assembly components were generated by rigidly docking trimeric components from their respective on-target design model into the off-target density maps. Off-target volumes exhibiting unexpected density regions of tetrameric component assembly were generated by running AlphaFold3 (*11*) with four copies of the corresponding construct sequence, followed by rigidly docking them into their corresponding volume maps. Models for I3-309 and I3-331 were additionally relaxed in ISOLDE with secondary structure restraints enabled.

### Biolayer interferometry

Octet Red96 and R8 Systems (Sartorius) were used for BLI. Antibodies 5J8 and FluA-20 were diluted in kinetics buffer (1×Phosphate-buffered saline (PBS), 0.5% serum bovine albumin and 0.01% Tween) to 0.02 mg/mL and protein samples were loaded at variable concentrations from 0.01-0.1 mg/mL. ProA Octet Biosensors (Sartorius) were used for loading antibodies and the following steps were performed at 25°C and 1000 rpm: 60 seconds buffer, 100 seconds loading, 30 seconds buffer, 100 seconds association, 100 seconds dissociation. Biosensors were regenerated with a total of 6 five second dips, alternating between 10 mM Glycine pH 2.5 and kinetics buffer.

### Cryo-EM sample preparation

Purified protein samples were prepared at concentrations of 0.3 mg/mL (O4-102), 0.5 mg/mL (I3-304 and O4-104), or 0.40 mg/mL (I3-326-MI15-TH) in 25 mM Tris, 150 mM NaCl. For I3-304, O4-102, and O4-104, 3 µL of sample was applied to glow-discharged Quantifoil R2.0/2.0 300-mesh grids with a 2 nm thin carbon support (16 s at 5 mA). For I3-326-MI15-TH, 2 µL of sample was applied to glow-discharged C-flat thick holey carbon grids. All grids were vitrified using a Vitrobot Mark IV operated at 22°C and 100% humidity. For Quantifoil grids, samples were incubated for 7.5 seconds prior to blotting with a blot time of 0.5 seconds and blot force of 0 or−1. For C-flat grids, samples were incubated for 7.5 seconds prior to blotting with a blot time of 6.5 seconds and blot force of 0. All grids were plunge-frozen into liquid ethane, clipped following standard procedures, and stored in liquid nitrogen prior to data collection.

### Cryo-EM data collection

Cryo-EM data were collected using SerialEM on Thermo Fisher Scientific transmission electron microscopes equipped with Gatan K3 direct electron detectors operated in counting mode. Data for O4-102 were collected on a Talos Glacios microscope operating at 200 kV, while data for I3-304, O4-102, O4-104, and I3-326 were collected on Titan Krios microscopes operating at 300 kV. For all datasets, images were acquired using beam-image shift with multi-hole acquisition schemes and fractionated electron exposures. For Talos Glacios data collection (O4-102), images were acquired with one shot per hole across nine holes per stage movement, with a total exposure of 28.8 e⁻/Å^2^ fractionated over 125 frames, yielding 234 movies. For Titan Krios data collection (I3-304, O4-102, and O4-104), microscopes were equipped with BioQuantum energy filters using a 15 eV slit width, and images were acquired with five shots per hole across nine holes per stage movement at a total exposure of approximately 50 e⁻/Å^2^ fractionated over 99 frames. In total, 6,302 movies were collected for I3-304, 6,375 movies for O4-102, and 11,590 movies for O4-104. Data for I3-326-MI15-TH were collected at the Janelia Research Campus on a Titan Krios microscope equipped with a Gatan energy filter using a 20 eV slit width. Images were recorded in super-resolution mode at a nominal magnification of 81,000×, corresponding to a physical pixel size of 1.061 Å. A total exposure of 50 e⁻/Å^2^ was fractionated over 75 frames per movie. Across all datasets, images were collected with nominal defocus ranges spanning −0.8 μm to −1.8 μm under stable cryogenic conditions.

### Cryo-EM data processing

All image processing was carried out in cryoSPARC and cryoSPARC live (Structura Biotechnology Inc.). Default parameters were used unless otherwise noted.

#### De novo icosahedral nanoparticle I3-304

Raw movies were subjected to Patch Motion Correction and Patch CTF estimation in cryoSPARC Live. Particles corresponding to the on-target icosahedral nanoparticle were identified using blob-based and template-based picking strategies and curated through multiple rounds of 2D classification. Following curation, 616,273 particles were retained and used for ab initio reconstruction and non-uniform refinement with imposed icosahedral symmetry. Iterative refinement incorporating per-particle defocus optimization, global CTF refinement, and per-particle scale-based subsetting reduced the dataset to 350,549 particles and yielded a final reconstruction at 2.40 Å global resolution (GSFSC, 0.143 cutoff). A corresponding asymmetric (C1) reconstruction at 2.90 Å resolution showed no evidence of symmetry breaking. In addition, a low-abundance off-target nanoparticle species (representing ∼1% of all particles) exhibiting D5 symmetry was identified. Due to its scarcity, 134 particles were initially identified by manual picking and used to generate 2D templates for automated particle picking. After curation and 2D classification, 8,592 particles were retained for 3D reconstruction. Ab initio reconstruction and non-uniform refinement were performed with imposed D5 symmetry, followed by global CTF refinement and reference-based motion correction. The final D5 refinement used 6,952 particles and reached a global resolution of 3.99 Å (GSFSC, 0.143 cutoff). A corresponding C1 reconstruction at 7.33 Å resolution showed no evidence of symmetry breaking. Final maps were sharpened using DeepEMHancer and assessed using orientation diagnostics and local resolution estimation. Reconstructions were deposited in the Electron Microscopy Database under accession numbers EMD-74566 (I3-304) and EMD-74499 (D5).

#### De novo octahedral nanoparticle O4-102

All image processing was performed in cryoSPARC and cryoSPARC Live (Structura Biotechnology Inc.) using default parameters unless otherwise noted. An initial dataset of 234 movies collected on a Glacios microscope was processed using Patch Motion Correction and Patch CTF estimation. Particles were identified using blob-based picking, curated through multiple rounds of 2D classification, and used for ab initio reconstruction. The dominant ab initio class exhibited clear octahedral symmetry and was refined homogeneously, yielding an initial reconstruction at 2.77 Å global resolution based on 46,945 particles. A higher-resolution dataset consisting of 6,375 movies collected on a Titan Krios microscope was processed in parallel using Patch Motion Correction and Patch CTF estimation. Template-based particle picking was performed using 2D class averages derived from the Glacios dataset, followed by iterative 2D classification and exposure curation. After particle curation, 510,009 particles were retained and subjected to heterogeneous refinement using the Glacios-derived reference, followed by non-uniform refinement. Iterative refinement incorporating per-particle defocus optimization, global CTF refinement, image-shift-based exposure grouping, and Ewald sphere curvature correction reduced the dataset to 320,100 particles and yielded a final reconstruction at 2.14 Å global resolution (GSFSC, 0.143 cutoff). Octahedral symmetry, observed in the initial ab initio reconstruction and consistent with the design model, was imposed during all symmetric refinements. To assess potential symmetry breaking, asymmetric and symmetry-relaxed non-uniform refinements were performed downstream, yielding reconstructions at 2.51 Å and 2.53 Å resolution, respectively, with no evidence of symmetry breaking. The final reconstruction was deposited in the Electron Microscopy Database under accession number EMD-74498.

#### De novo octahedral nanoparticle O4-104

All image processing was performed in cryoSPARC and cryoSPARC Live (Structura Biotechnology Inc.) using default parameters unless otherwise noted. Raw movies were subjected to Patch Motion Correction and Patch CTF estimation in cryoSPARC Live. Particles were identified using blob-based picking strategies, extracted, and curated through exposure curation, micrograph denoising, and multiple rounds of 2D classification to remove false positives, ice contamination, and poorly aligned particles. Following particle curation, 570,865 particles were retained and used for ab initio reconstruction with C1 symmetry. Subsequent non-uniform refinement with imposed octahedral symmetry and per-particle defocus optimization yielded an initial reconstruction at 2.31 Å global resolution. Particles were further subset based on per-particle scale contribution, resulting in a final particle set of 402,797 particles. These particles were refined using iterative non-uniform refinement incorporating image-shift-based exposure grouping, global CTF refinement, and Ewald sphere curvature correction, yielding a final reconstruction at 2.14 Å global resolution (GSFSC, 0.143 cutoff). The final reconstruction was deposited in the Electron Microscopy Database under accession number EMD-74530.

#### De novo icosahedral nanoparticle I3-326-MI15-TH

All image processing was performed in cryoSPARC (Structura Biotechnology Inc.) using default parameters unless otherwise noted. Raw movies collected at Janelia Research Campus were subjected to Patch Motion Correction and Patch CTF estimation, followed by manual exposure curation to remove low-quality micrographs. An initial subset of 164 micrographs was manually picked to generate reference 2D class averages for template-based particle picking across the curated dataset. Following particle curation and multiple rounds of 2D classification, 38,017 particles were retained for 3D reconstruction. An initial ab initio reconstruction was generated with imposed icosahedral symmetry and refined using non-uniform refinement with per-particle defocus optimization, yielding a reconstruction at 3.11 Å global resolution. A masks corresponding to the nanoparticle core was generated using volume segmentation and the corresponding design model, respectively. Symmetry expansion followed by local refinement yielded a reconstruction for the nanoparticle core, refined with icosahedral symmetry using 38,017 particles, with a final resolution of 3.10 Å (GSFSC, 0.143 cutoff). The final reconstruction was deposited in the Electron Microscopy Database under accession number EMD-74706.

### Cryo-EM model building, refinement, and validation

For all structures, the corresponding computational design models were used as initial references for atomic model building. Design models were docked into the final sharpened cryo-EM density maps using UCSF Chimera (*42*) or ChimeraX (*43*). Where local deviations from the design were observed, individual subunits or asymmetric units were extracted and manually rebuilt in Coot (*44–46*). For all on-target icosahedral and octahedral reconstructions, a single chain was refined prior to symmetrization, whereas for the D5 off-target reconstruction, three chains corresponding to a trimeric asymmetric unit were refined together. Atomic models were iteratively refined using a combination of automated and manual approaches. Initial relaxation was performed using NAMDINATOR (*47*), followed by manual inspection and correction in Coot (*44–46*) and ISOLDE (*48*). Multiple rounds of relaxation and real-space refinement were carried out until convergence of model geometry and agreement with the density were achieved. Final real-space refinement was performed in Phenix (*49*). Refined models were re-symmetrized in ChimeraX (*43*), with limited additional refinement where necessary. Model quality was assessed using MolProbity (*50*). Figures were generated using UCSF Chimera (*42*) or ChimeraX (*43*). Final structures were deposited in the Protein Data Bank under the following accession numbers: I3-304 (9ZOL, 9ZQI), O4-102 (9ZOJ), O4-104 (9ZPM), and I3-326-MI15-TH (9ZSD).

### Cryo-EM water molecule placement and validation

Initial water molecules were placed using Phenix.douse on the octahedral symmetry refined maps with a stricter map_threshold_scale of 0.7 to limit placement to higher-confidence peaks. All Douse-generated waters were then evaluated in ChimeraX (*43*) using CheckWaters, and any waters lacking clear density support or plausible hydrogen bonding were removed. Manual inspection and rebuilding of the remaining waters were carried out in Coot (*44–46*) around a single selected chain. After manual correction and the addition of any justified waters, this edited chain and its associated waters were propagated to all sixteen subunits in ChimeraX (*43*) using the octahedral symmetry operators to produce a complete solvent model for the full particle. The propagated models were refined in ISOLDE (*48*), allowing the waters to relax into favorable positions that matched the surrounding density. The final coordinates were then run through Phenix (*49*) real-space refinement using only minimization, with the maximum number of iterations set to 100 and the number of macrocycles set to 1.

### Crystallographic data collection and structure determination

Crystallization experiments were conducted using the sitting drop vapor diffusion method. Initial crystallization trials were set up in 200 nL drops using the 96-well plate format at 20°C. Crystallization plates were set up using a Mosquito LCP from SPT Labtech, then imaged using UVEX microscopes and UVEX PS-256 from JAN Scientific. Diffraction quality crystals formed in 0.1 M Na HEPES, 1 M sodium acetate in 2 weeks for T2-71.

Diffraction data was collected at the National Synchrotron Light Source II on beamline 17-ID-2. X-ray intensities and data reduction were evaluated and integrated using XDS (*51*) and merged/scaled using Pointless/Aimless in the CCP4 program suite (*52*). Structure determination and refinement starting phases were obtained by molecular replacement using Phaser (*53*) using the design model for the structures. Following molecular replacement, the models were improved using phenix.autobuild with rebuild-in-place to false, and using simulated annealing. Structures were refined in Phenix (*49*). Model building was performed using Coot (*44*). The final model was evaluated using MolProbity (*50*). Data deposition, atomic coordinates, and structure factors reported in this paper have been deposited in the Protein Data Bank (PDB), http://www.rcsb.org/ with accession code 9ZV3.

### In vivo study ethics statement

Animal experiments were conducted in accordance with the NIH Guide for the Care and Use of Laboratory Animals and under approval by the University of Washington’s Institutional Animal Care and Use Committee (IACUC protocol 4470-01). The University of Washington’s animal use program is accredited by the Association for Assessment and Accreditation of Laboratory Animal Care (Reference Assurance: #000523). Mice were maintained in a specific-pathogen free facility within the Department of Comparative Medicine where housing is temperature and humidity-controlled, standard light/dark cycles are used, and access to food and water is provided *ad libitum*.

### Immunization

Female BALB/cAnNHsd mice were ordered from Envigo at 7 weeks of age. Mice were immunized at 8 weeks of age subcutaneously in the inguinal region with 1 µg of low-endotoxin protein in 100 µL volume delivered as a 1:1 mixture with AddaVax adjuvant (InvivoGen vac-adx-10) at weeks 0 and 4. At week 6, terminal bleeds were performed via cardiac puncture and whole blood was collected in serum separator tubes (BD #365967). After allowing the blood to rest at room temperature for 30 minutes for coagulation, tubes were spun at 2,000 *g* for 10 minutes to collect the serum, which was stored at −80°C until use.

### Enzyme-linked immunosorbent assay (ELISA)

Full-length wild-type HA ectodomain fused to the trimeric nanoparticle component I53_dn5B was plated on 96-well half-area microplates (Corning) at 0.02 mg/mL with 50 µL per well and incubated for 18 hours at 4°C. After incubation, 100 µL of blocking buffer (TBST: 25 mM Tris pH 8.0, 150 mM NaCl, 0.05% (v/v) Tween 20, and 5% Nonfat milk) was added to each well and incubated once more at room temperature for 1 hour. Plates were then washed three times with TBST and 50 µL of serum dilutions were added to each well. Three-fold serum dilutions were made for all animals in every group, starting with a 1:100 serum dilution. Following 45 minutes of incubation at room temperature, plates were again washed three times with TBST and 50 µL of peroxidase AffiniPure goat anti-mouse IgG (Fcγ fragment specific, Jackson Immunoresearch) was added to each well at a 1:2,000 dilution before a final 30 minute, room temperature incubation. Plates were washed one more time and 50 µL of TMB (3,30,50,5-tetramethylbenzidine, SeraCare) was added for 5 minutes, followed by quenching with 50 µL per well of 1 N HCl. Reading at 450 nm absorbance was done on a Epoch plate reader (BioTek).

### Hemagglutination inhibition (HAI)

Serum was inactivated using receptor-destroying enzyme (RDE) II (Seiken) in PBS at a 3:1 ratio of RDE to serum for 18 hours at 37°C, followed by 40 minutes at 56°C. For each animal, 20 µL inactivated serum was mixed with 30 µL PBS and serially diluted 2-fold in V-bottom plates at 25 µL per well. 25 µL HA-ferritin nanoparticles was added to each well and incubated at room temperature for 30 minutes. Turkey red blood cells (Lampire) were prepared by centrifugation at 500 *g* for 5 minutes and resuspension of the pellet to 1% in PBS. After the incubation step, 50 µL of the prepared red blood cells were added to each well and incubated for at least 2 hours prior to recording HAI titers, reported as the last dilution at which agglutination was prevented by serum.

### Influenza microneutralization assay

The A/Michigan/45/2015(H1N1) virus was prepared as a replication-restricted reporter (R3) DPB1 virus as previously described (*54*). Briefly, the virus encodes an engineered PB1 segment in which the reporter gene TdKatushka2, fused to a C-terminal nuclear localization signal, was inserted between the PB1 genomic packaging sequences of A/WSN/1933 (H1N1), replacing most of the PB1 coding region (Genbank MW298217.1). The internal genes (PB2, PA, NP, M, and NS) were derived from A/WSN/1933 (H1N1), while the HA (Genbank MW298181.1) and NA (Genbank MW298248.1) segments were from A/Michigan/45/2015 (H1N1). The R3 DPB1 virus was rescued by reverse genetics in 293 cells stably expressing PB1 of A/WSN/1933 (H1N1) and propagated in MDCK-SIAT1 (Millipore Sigma) cells constitutively expressing PB1 of A/WSN/1933 (H1N1) (MDCK-SIAT1-PB1 cell line) in the presence of TPCK-trypsin (Millipore Sigma).

For the serum microneutralization assay, mouse serum samples were treated with RDE by mixing one volume of serum with three volumes of RDE and incubating overnight at 37°C, followed by heat inactivation at 56°C. Three replicates of four-fold serial dilutions of RDE-treated sera were incubated with an equal volume of pre-titrated virus for 1 h at 37°C in a 5% CO_2_ humidified atmosphere. MDCK-SIAT1-PB1 cells were then added, and the serum-virus-cell mixtures were transferred to 384-well plates and incubated overnight at 37°C in 5% CO_2_ humidified atmosphere.

The following day, fluorescent objects were automatically detected and quantified using a Celigo image cytometer (Revvity) with a customized red filter to detect TdKatushka2 fluorescence (*54*). Percent neutralization was calculated by constraining the virus-only control as 0% neutralization and the cell-only control as 100% neutralization. Neutralization curves were generated by plotting percent neutralization against serum concentration, and 80% inhibitory concentrations (IC₈₀) were estimated using a sigmoidal four-parameter logistic (4PL) fit in Prism (GraphPad).

## Supplementary Data

**Figure S1.**
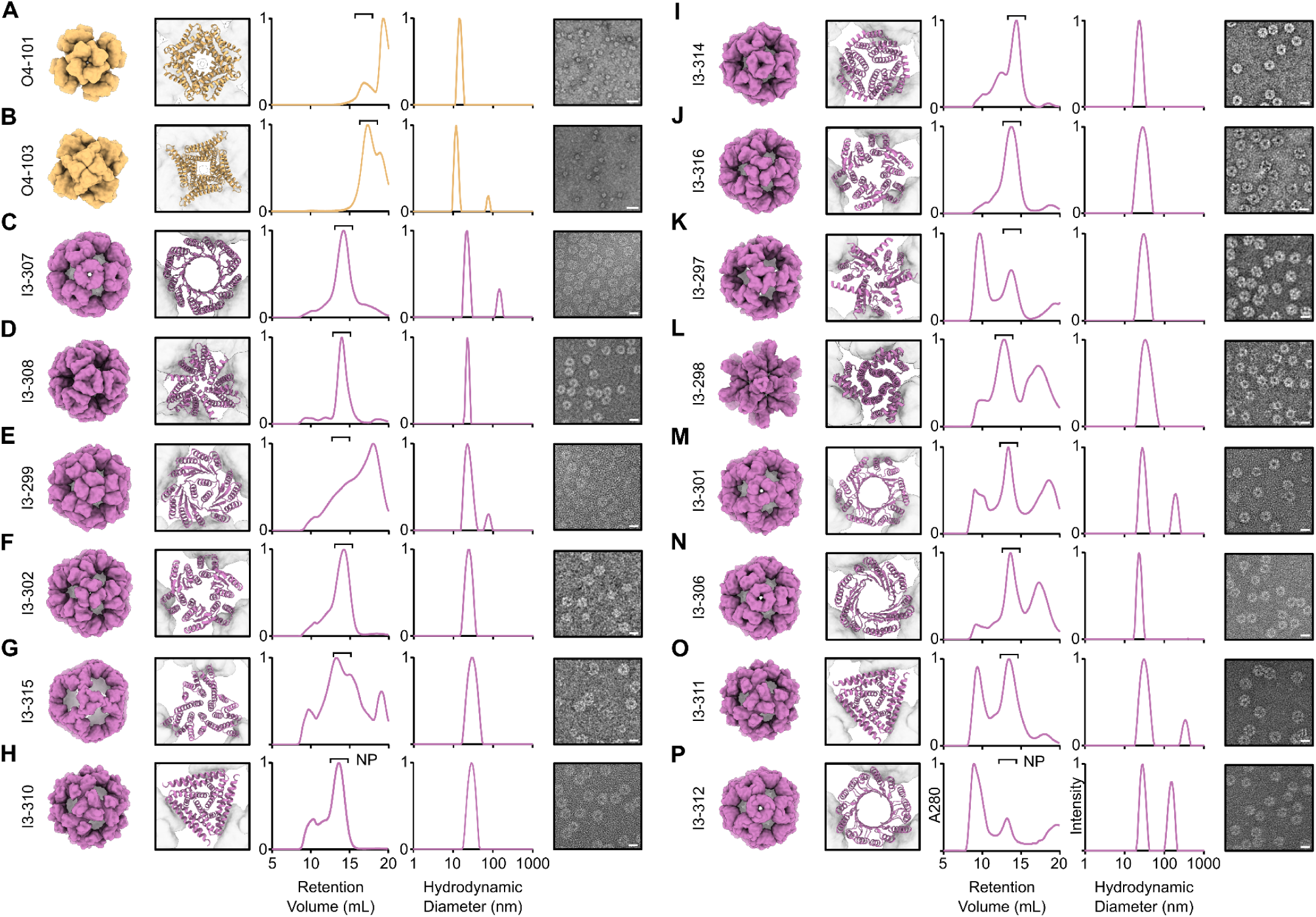
Characterization of de novo nanoparticles generated with unconditional diffusion of components. *From left to right:* Design models, SEC, DLS (intensity), and negatively stained electron micrographs for (**A**) O4-101, (**B**) O4-103, (**C**) I3-307, (**D**) I3-308, (**E**) I3-299, (**F**) I3-302, (**G**) I3-315, (**H**) I3-310, (**I**) I3-314, (**J**) I3-316, (**K**) I3-297, (**L**) I3-298, (**M**) I3-301, (**N**) I3-306, (**O**) I3-311, and (**P**) I3-312. The design model of the nanoparticle is shown at left and a cropped image of the design model for the diffused oligomer in the context of the nanoparticle is shown at right. nsEM scale bars = 20 nm.

**Figure S2.**
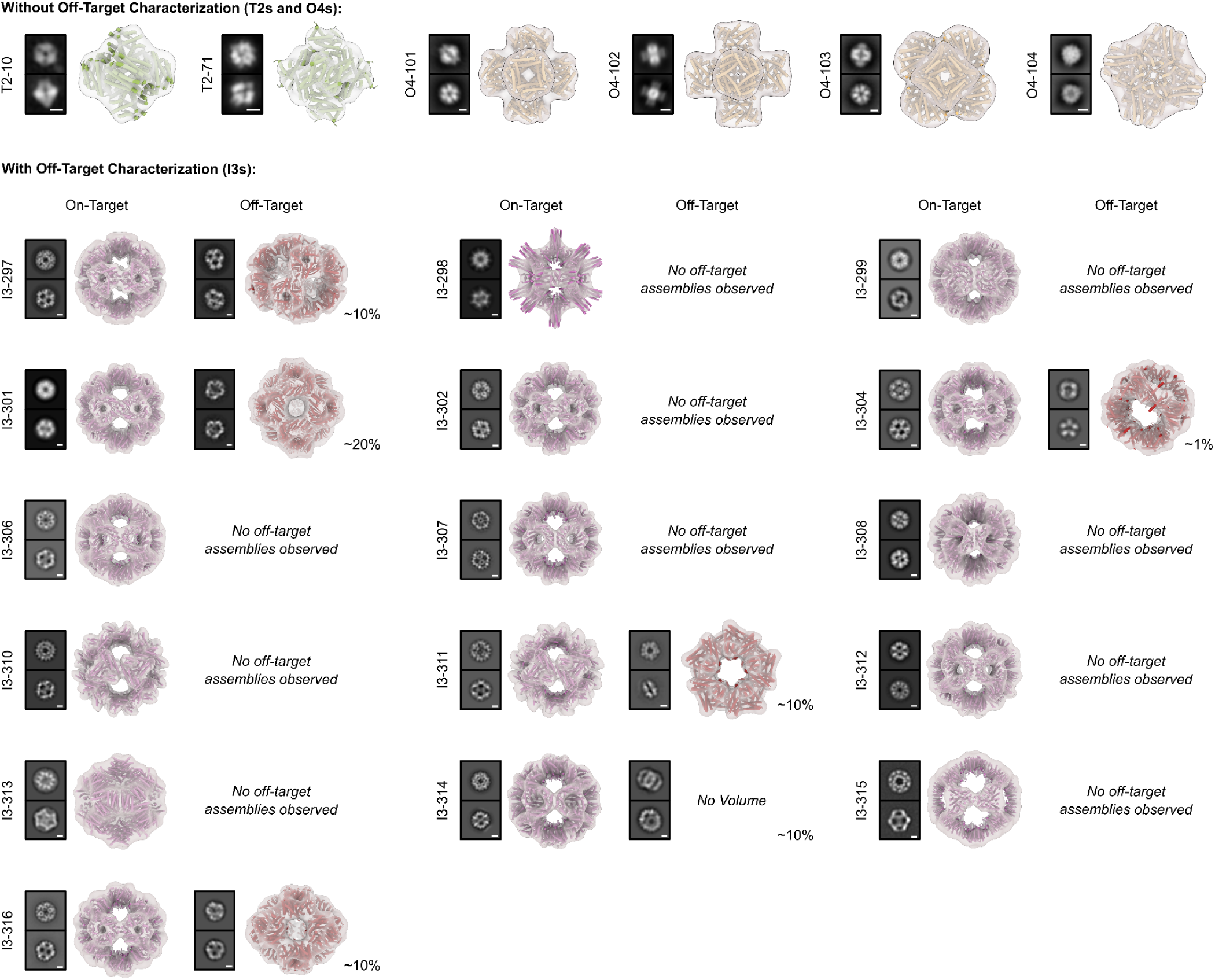
nsEM analysis of unconditionally diffused nanoparticles. *Left,* 2D class averages and *right,* 3D reconstructions of on-target nanoparticles and off-target assemblies (when present). Off-target species were only analyzed for successfully designed icosahedral nanoparticles. Reconstructions show the map in transparent grey with the model of the protein nanoparticle shown underneath as a cartoon in pink (icosahedral), yellow (octahedral), green (tetrahedral), or red (off-target). Relative abundances of observed off-target species are approximated next to each species. Class averages for nanoparticles shown in Figure 2 are replicated here for clarity. Scale bars = 5 nm.

**Figure S3.**
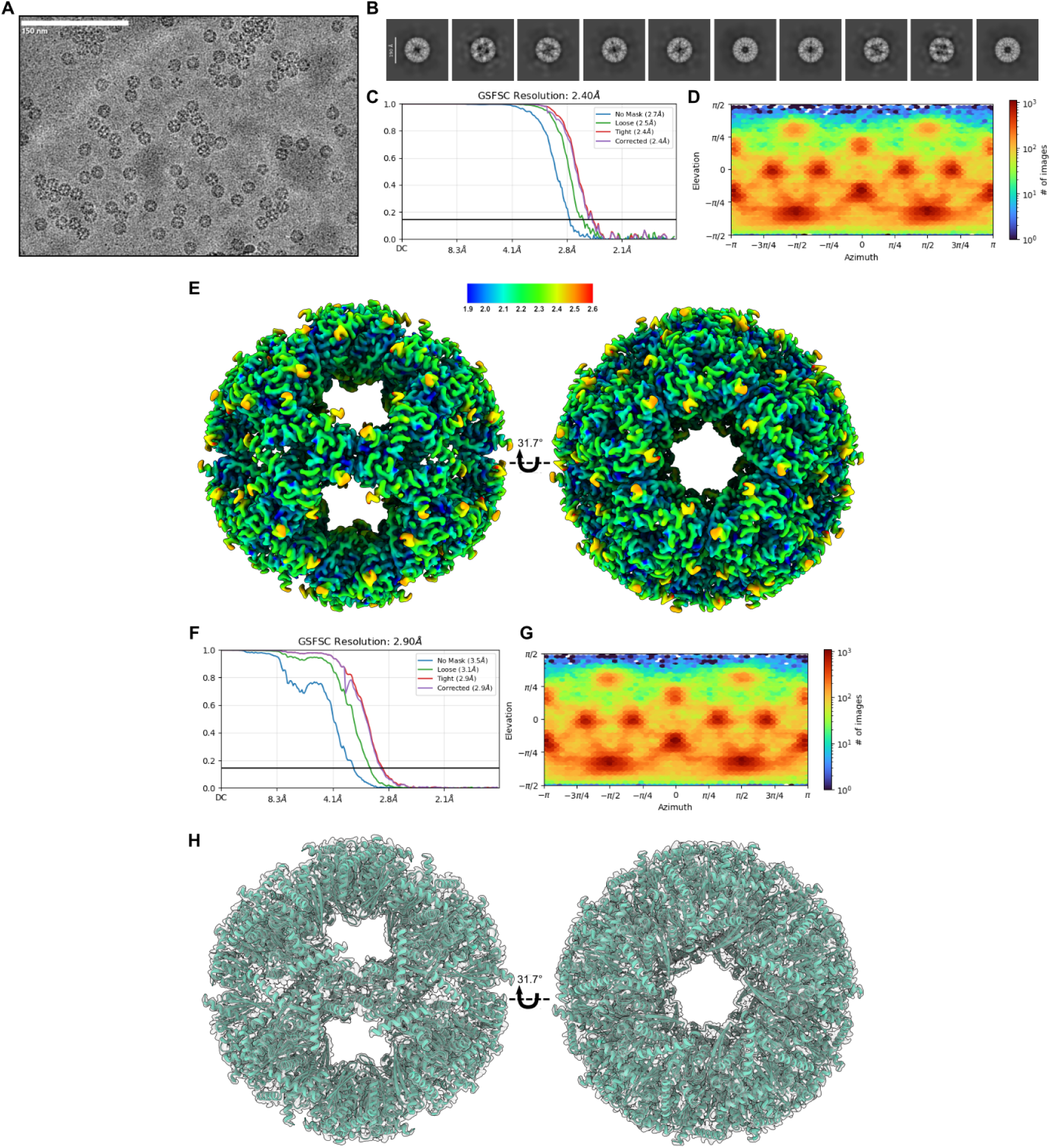
Cryo-EM analysis of de novo designed octahedral nanoparticle I3-304. (**A**) Representative cryo-EM micrograph. (**B**) Representative 2D class averages of the dominant on-target I3-304 nanoparticle. (**C**) Global gold-standard FSC from non-uniform refinement with icosahedral symmetry applied, yielding a resolution of 2.90 Å at the 0.143 cutoff. (**D**) Orientational distribution plot demonstrating full angular sampling. (**E**) Local resolution map (0.143 FSC cutoff). (**F**) Global gold-standard FSC from non-uniform refinement with C1 symmetry applied. (**G**) Orientational distribution plot from C1 refinement. (**H**) Icosahedral atomic model docked into the C1 map, showing no detectable deviations from imposed symmetry.

**Figure S4.**
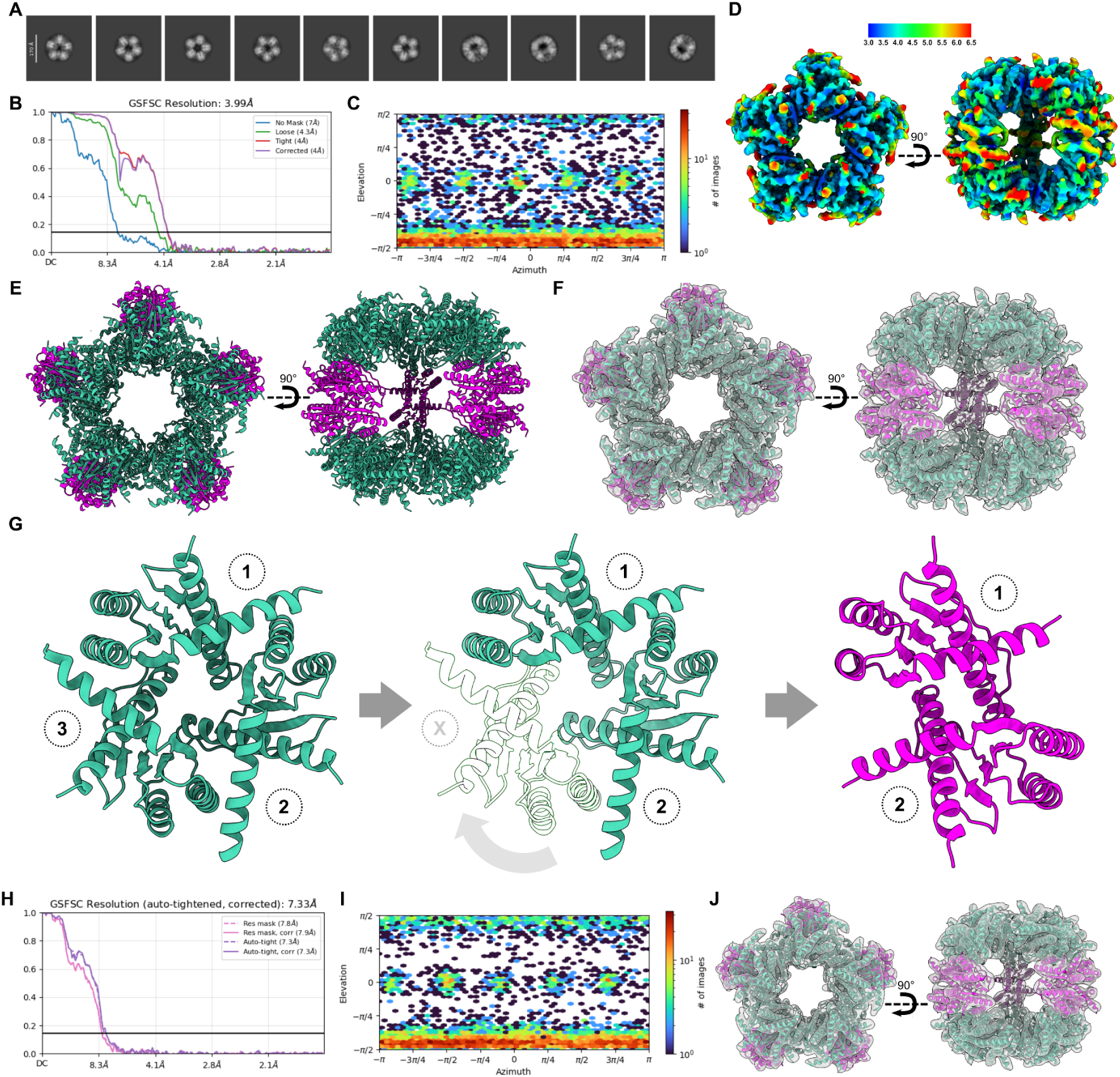
Cryo-EM analysis of the I3-304 off-target D5 nanoparticle assembly. (**A**) Representative 2D class averages. (**B**) Global gold-standard Fourier shell correlation (GSFSC) from non-uniform refinement with D5 symmetry, yielding a resolution of 3.99 Å at the 0.143 cutoff. (**C**) Orientational distribution plot. (**D**) Local resolution map (0.143 FSC cutoff). (**E**) Atomic model of the off-target D5 assembly. The assembly comprises 40 subunits, arranged as two sets of five trimers (green) connected by five off-target dimers (pink). (**F**) Atomic model docked into the 3.99 Å D5-symmetrized cryo-EM density map, demonstrating high model-to-map agreement. (**G**) Structural analysis of the off-target dimer observed at the dihedral two-fold axes. *Left,* the on-target trimeric building block; *middle,* elimination of subunit X of the trimer and rotation of subunit 2 leads to *right,* the off-target dimer (pink), which forms a C2-symmetric dimer in the absence of the third chain. (**H**) Global gold-standard Fourier shell correlation (GSFSC) from non-uniform refinement performed in C1 symmetry, yielding a resolution of 7.33 Å at the 0.143 cutoff. (**I**) Orientational distribution plot for the C1 refinement. (**J**) D5 atomic model rigid-body docked into the C1 cryo-EM density map, showing no evidence of symmetry deviations and confirming the D5 architecture of the off-target assembly.

**Figure S5.**
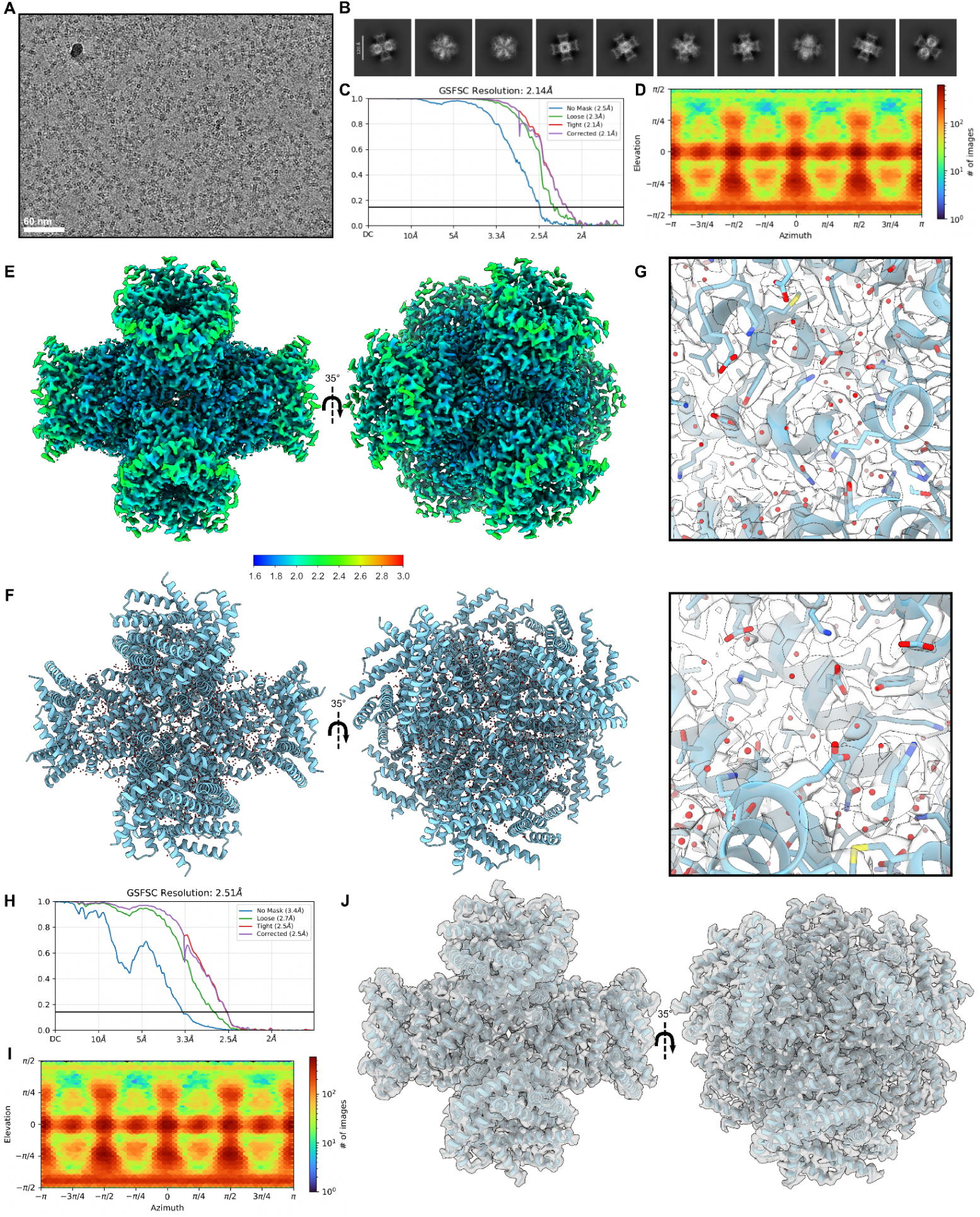
Cryo-EM analysis of de novo designed octahedral nanoparticle O4-102. (**A**) Representative raw micrograph showing well-defined particles with some evidence of flocculation and aggregation. Scale bar, 60 nm. (**B**) 10 representative two-dimensional (2D) class averages displaying multiple particle orientations. (**C**) Global gold-standard Fourier shell correlation (GSFSC) curve indicating an estimated global resolution of 2.14 Å. Orientational distribution plot demonstrating full angular sampling. (**E**) Local resolution map colored by resolution and shown along the 2- and 3-fold axes of symmetry. (**F**) Atomic model of O4-102 viewed along the 2- and 3-fold axes of symmetry, with ordered water molecules shown. (**G**) Representative map-to-model fit, shown from two angles. (**H-J**) Asymmetric refinement demonstrating a lack of symmetry-breaking features. (**H**) GSFSC curve indicating an estimated global resolution of 2.51 Å. (**I**) Orientational distribution plot demonstrating full angular sampling. (**J**) Atomic model rigid-body docked inside the unsharpened C1-refined cryo-EM density map.

**Figure S6.**
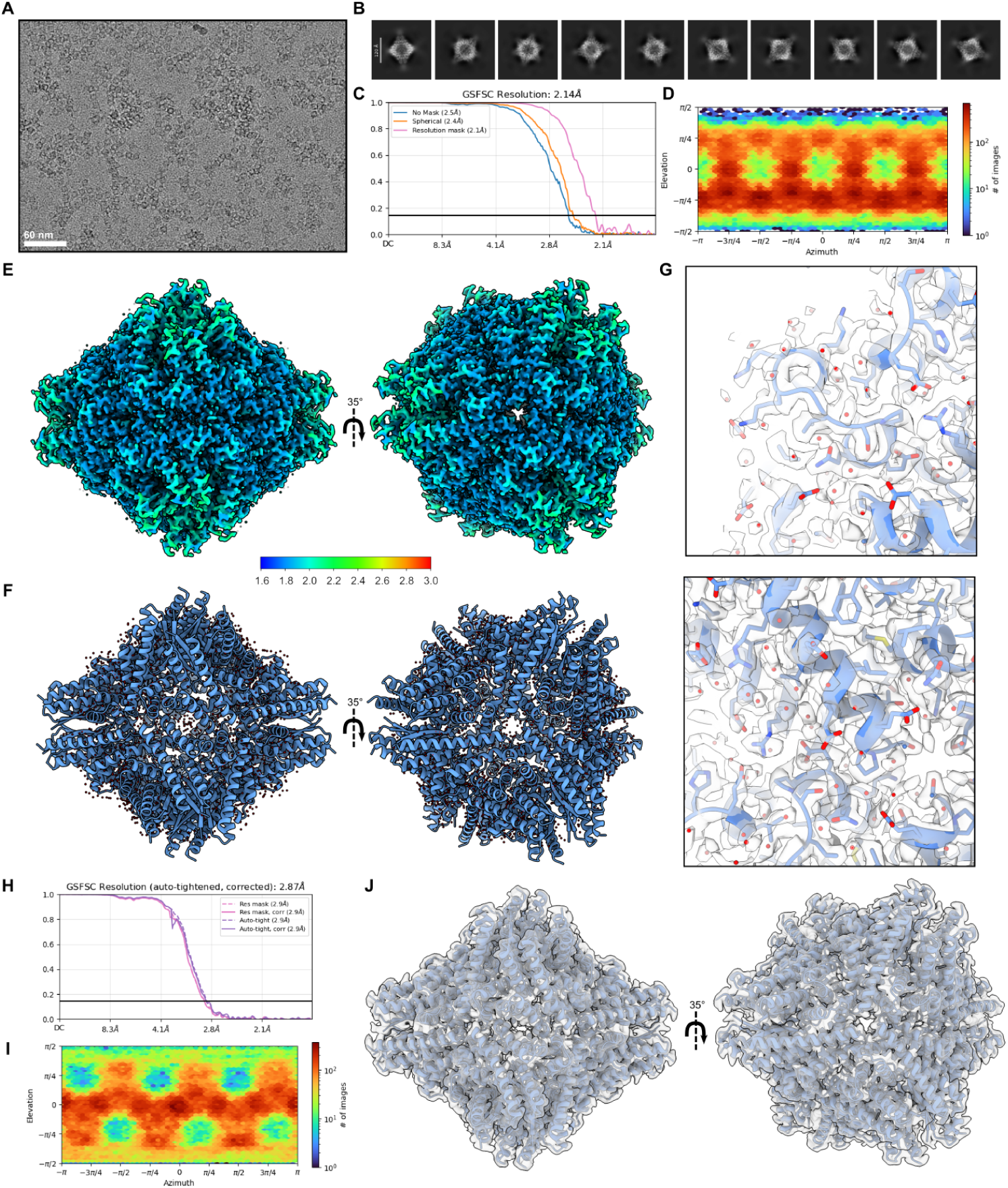
Cryo-EM analysis of de novo designed octahedral nanoparticle O4-104. (**A**) Representative raw micrograph showing well-defined particles with some evidence of flocculation and aggregation. Scale bar, 60 nm. (**B**) 10 representative two-dimensional (2D) class averages displaying multiple particle orientations. (**C**) Global gold-standard Fourier shell correlation (GSFSC) curve indicating an estimated global resolution of 2.14 Å. (**D**) Orientational distribution plot demonstrating full angular sampling. (**E**) Local resolution map colored by resolution and shown along the 2- and 3-fold axes of symmetry. (**F**) Atomic model of O4-104 viewed along the 2- and 3-fold axes of symmetry, with ordered water molecules shown. (**G**) Representative map-to-model fit, shown from two angles. (**H-J)** asymmetric refinement demonstrating a lack of symmetry breaking features. (**H**) GSFSC curve indicating an estimated global resolution of 2.87 Å. (**I**) Orientational distribution plot demonstrating full angular sampling. (**J**) Atomic model rigid-body docked inside the unsharpened C1-refined cryo-EM density map.

**Figure S7.**
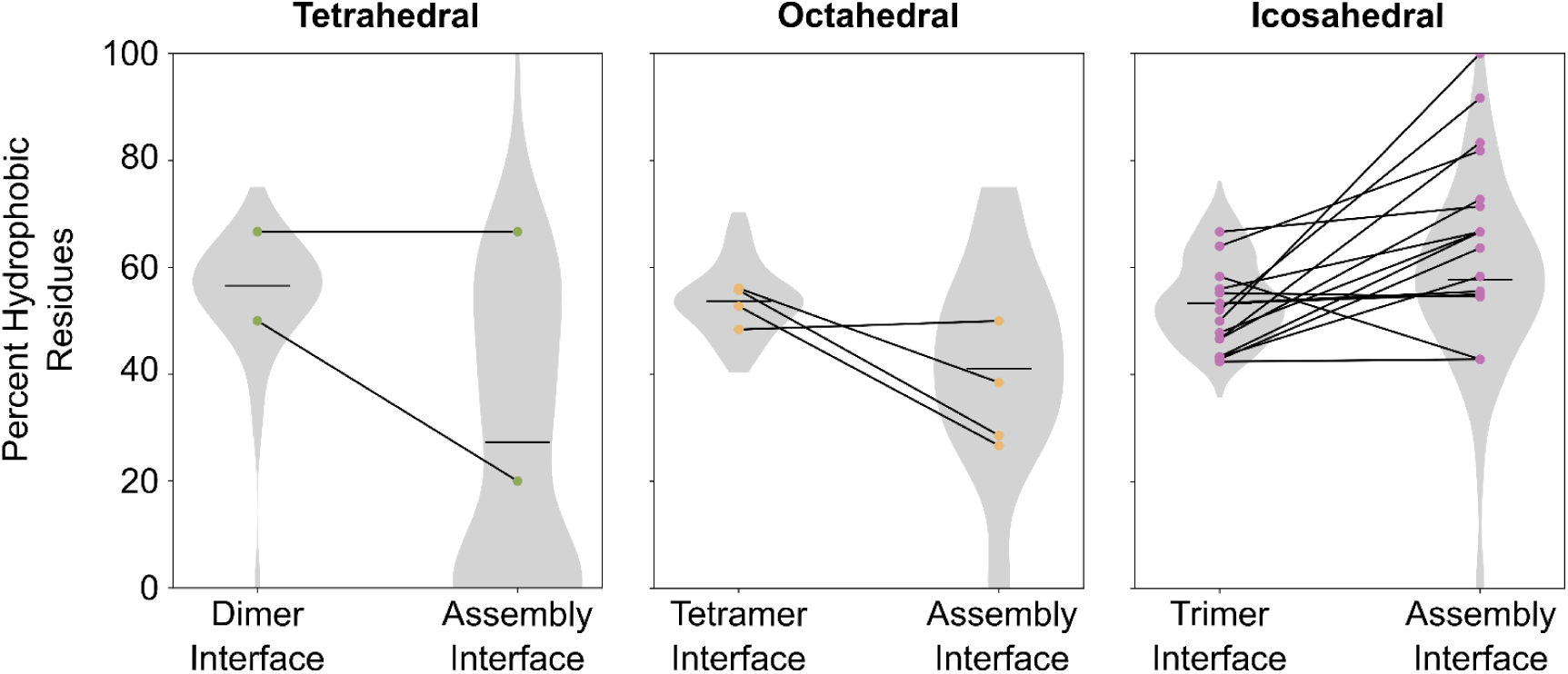
Analysis of hydrophobic residues at interfaces for the tetrahedral, octahedral, and icosahedral nanoparticles selected for experimental characterization. Violin plots showing the distribution of the percent hydrophobic residues (A, V, L, I, M, F, W, P, Y) found at the oligomer and nanoparticle assembly interfaces for designs that were ordered for experimental validation (tetrahedral, n=84; octahedral, n=56; icosahedral, n=166). Assembly interfaces were defined as the set of residues including Cɑ atoms within 8 Å of a Cɑ on another chain in the design model that was not part of the same dimeric, trimeric, or tetrameric building block. Points correspond to nanoparticles that were experimentally confirmed to successfully assemble to the target architecture.

**Figure S8.**
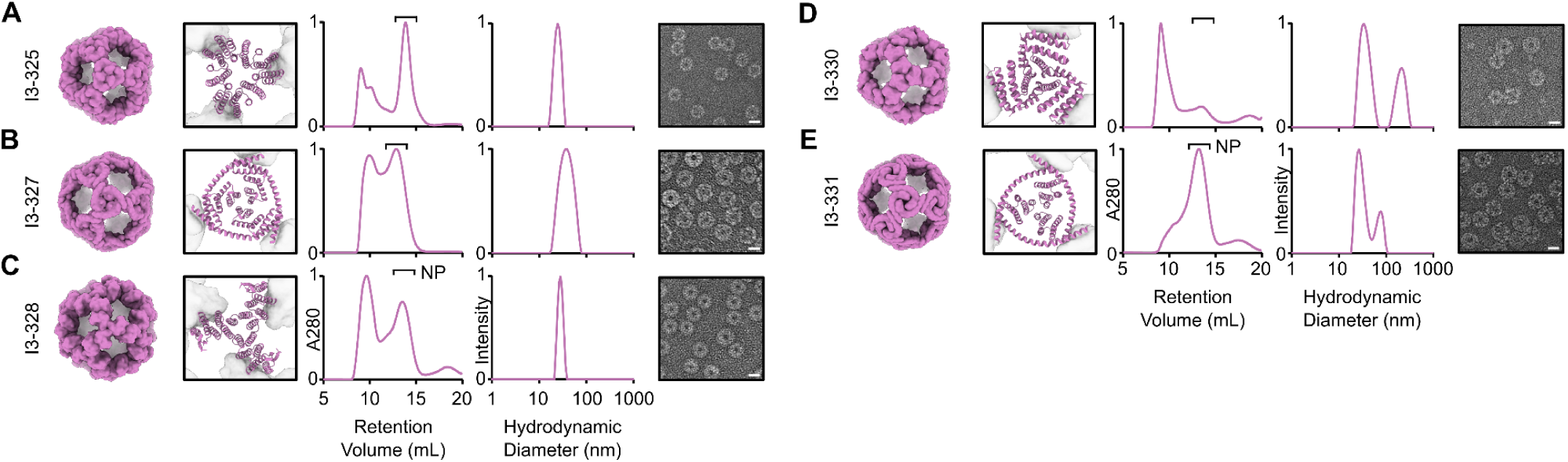
Characterization for additional motif-scaffolded icosahedral nanoparticles. *From left to right:* Design models, SEC, DLS, and negatively stained electron micrographs for (**A**) I3-325, (**B**) I3-327, (**C**) I3-328, (**D**) I3-330, and (**E**) I3-331. The design model of the nanoparticle is shown at left and a cropped image of the design model for the diffused oligomer in the context of the nanoparticle is shown at right. nsEM scale bars = 20 nm.

**Figure S9.**
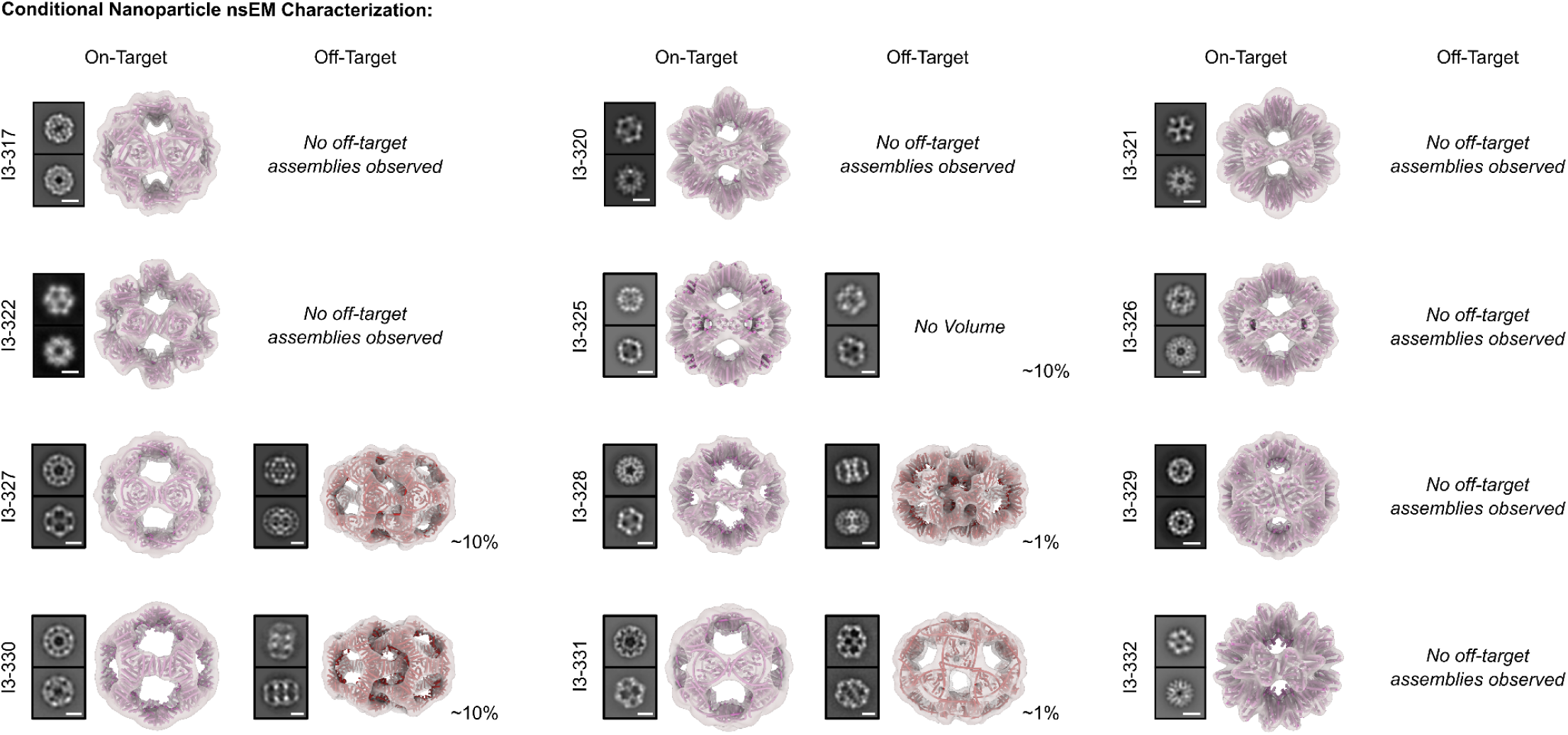
nsEM analysis of antigen-tailored icosahedral nanoparticles. *Left,* 2D class averages and *right,* 3D reconstructions of on-target nanoparticles and off-target assemblies (when present). Reconstructions show the map in transparent gray with the model of the protein nanoparticle shown underneath as a cartoon in pink or red. Relative abundances of observed off-target species are approximated next to each species. Class averages for nanoparticles shown in Figure 4 are replicated here for clarity. Scale bars = 10 nm.

**Figure S10.**
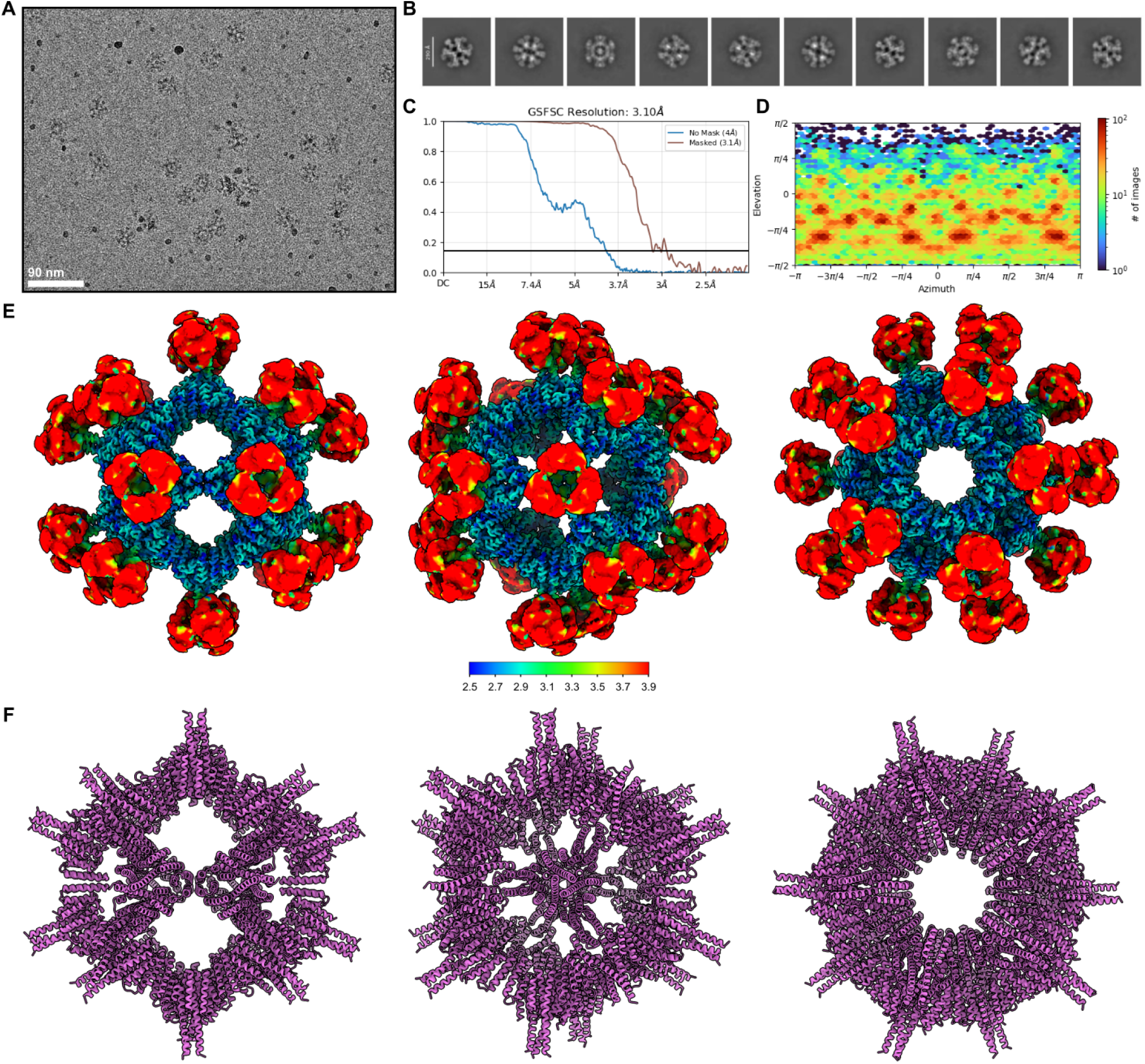
Cryo-EM analysis of de novo designed icosahedral nanoparticle I3-326 displaying HA trihead antigen. (**A**) Representative raw micrograph showing well-defined particles with no evidence of flocculation or aggregation. Scale bar, 90 nm. (**B**) 10 representative two-dimensional (2D) class averages displaying multiple particle orientations. (**C**) Global gold-standard Fourier shell correlation (GSFSC) curve indicating an estimated global resolution of 3.10 Å. (**D**) Orientational distribution plot demonstrating near-full angular sampling. (**E**) Local resolution map colored by resolution and shown along the three principal symmetry axes. (**F**) Cryo-EM density and atomic model of I3-326 viewed along the three major symmetry axes.

**Figure S11.**
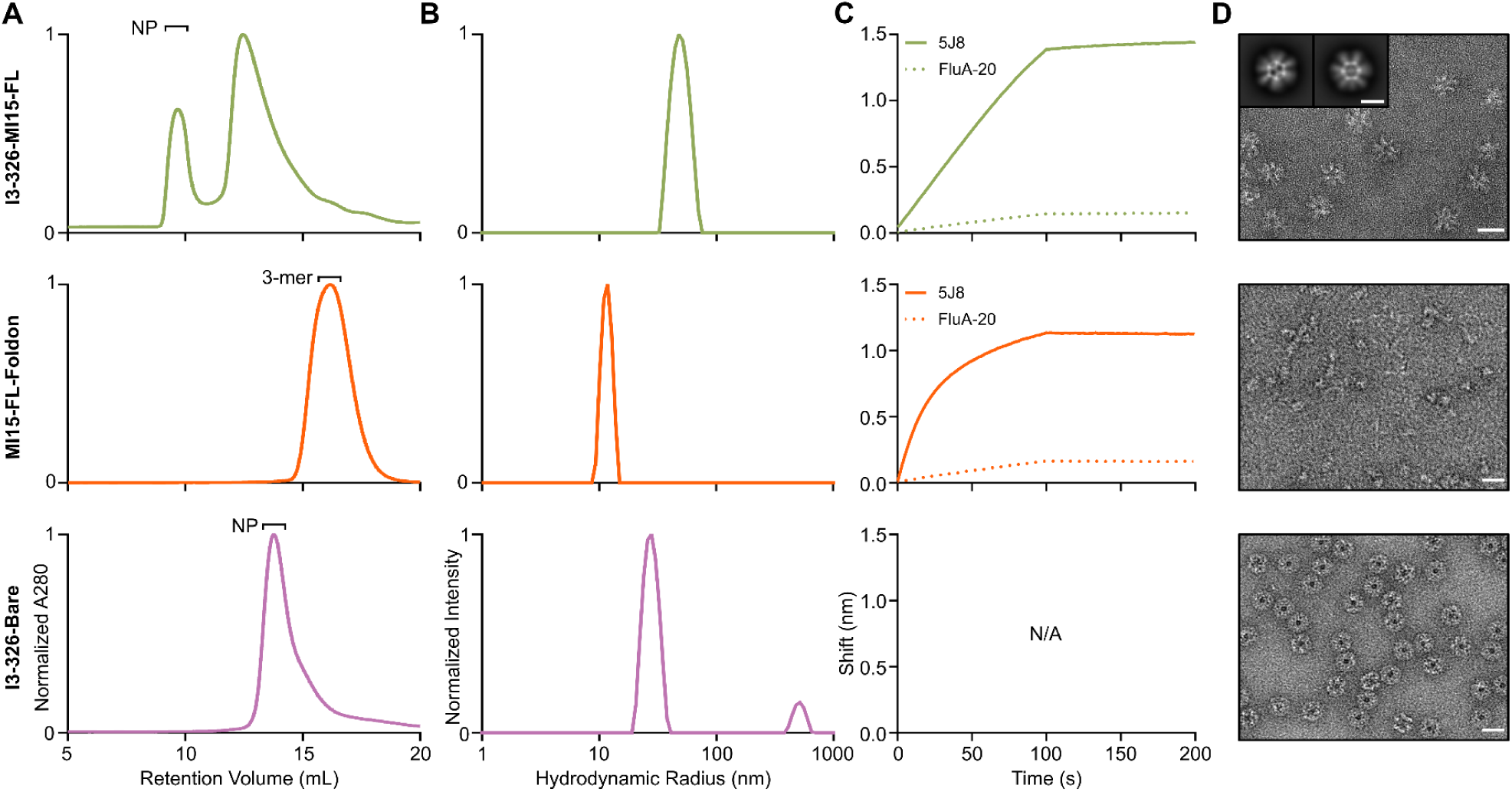
Characterization of proteins used in immunogenicity study. (**A**) SEC, (**B**) DLS, (**C**) BLI, and (**D**) nsEM for the proteins used in the immunogenicity study. SEC and DLS plots are normalized and fractions purified for further characterization are indicated (“NP”, nanoparticle; “3-mer”, trimer). 2D class averages are shown for I3-326-MI15-FL. Scale bars = 50 nm, 20 nm, and 20 nm for I3-326-MI15-FL, MI15-FL-Foldon, and I3-326-Bare, respectively. Scale bar = 25 nm for I3-326-MI15-FL 2D class averages.

**Figure S12.**
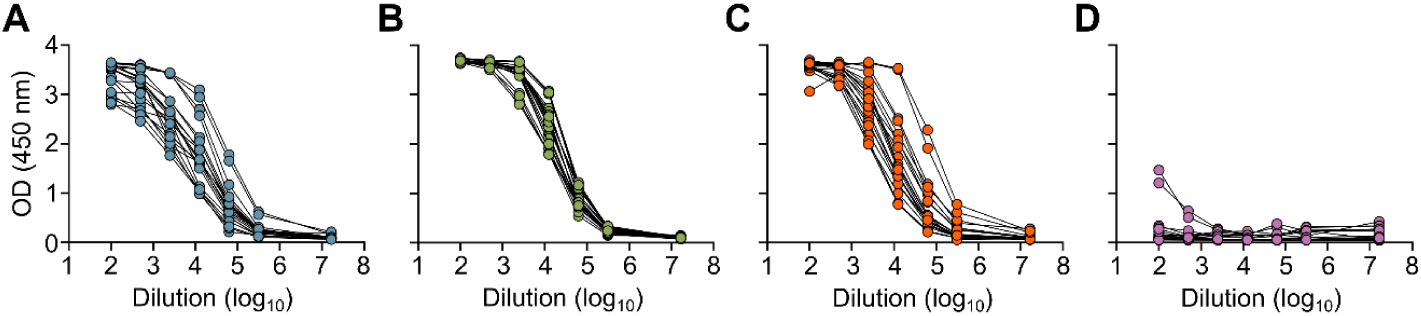
Raw ELISA data and fits used to determine titers shown in Figure 5. The ELISA curves showing week six serum IgG binding to full-length HA ectodomain trimer (A/Michigan/45/2015) for (**A**) I3-326-MI15-TH, (**B**) I3-326-MI15-FL, (**C**) MI15-FL-Foldon, and (**D**) I3-326-Bare.

**Table S1.**
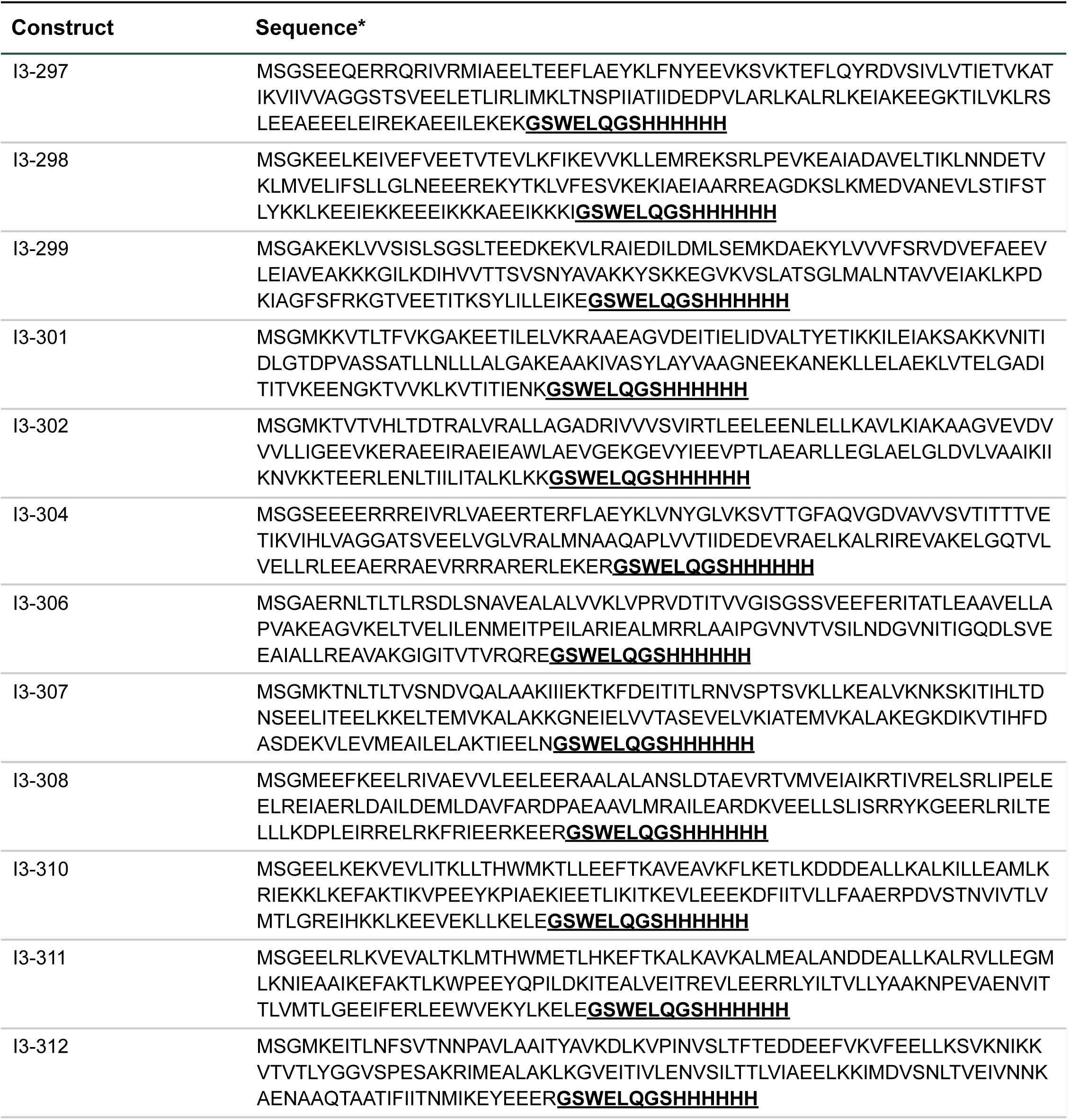
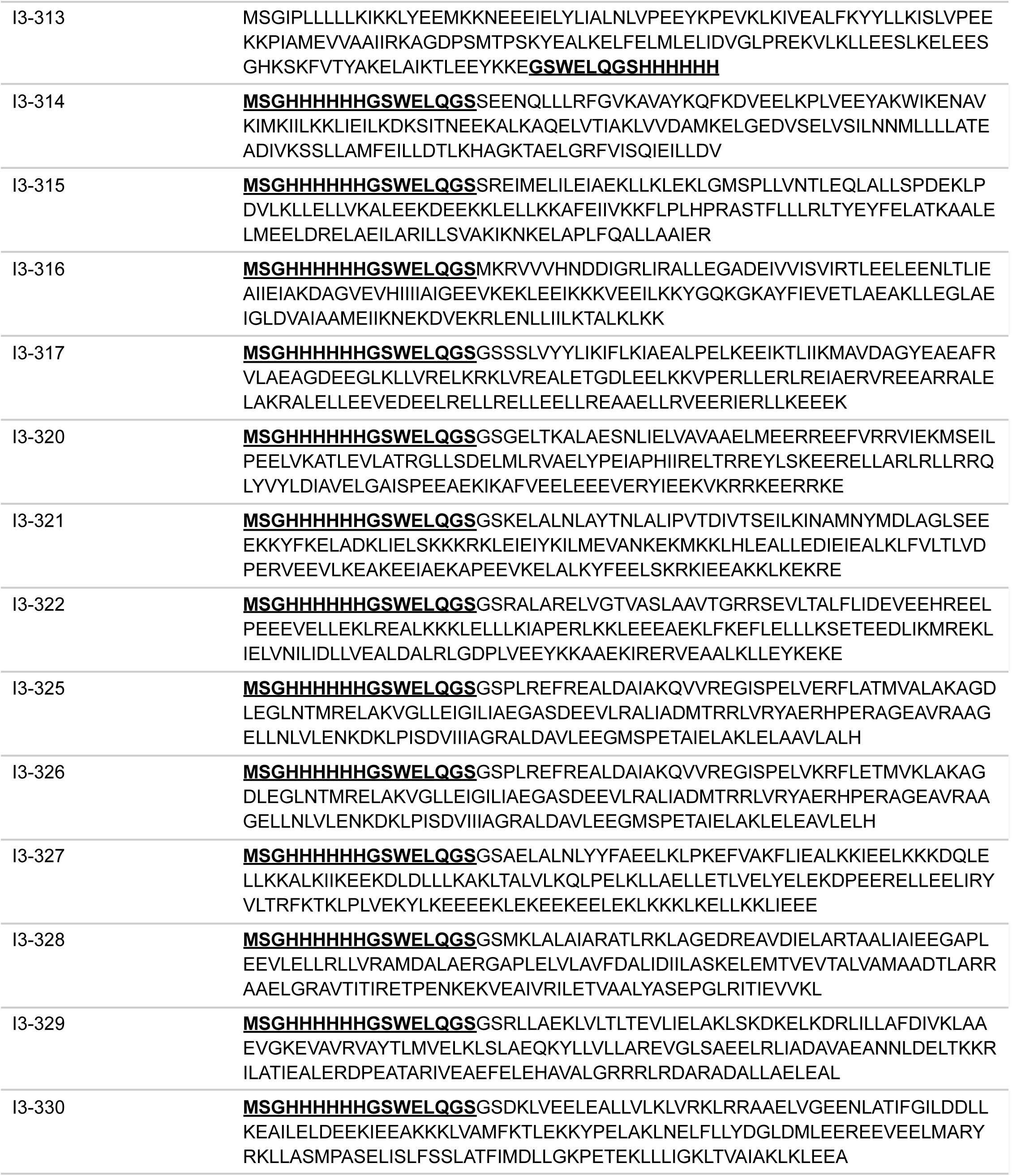
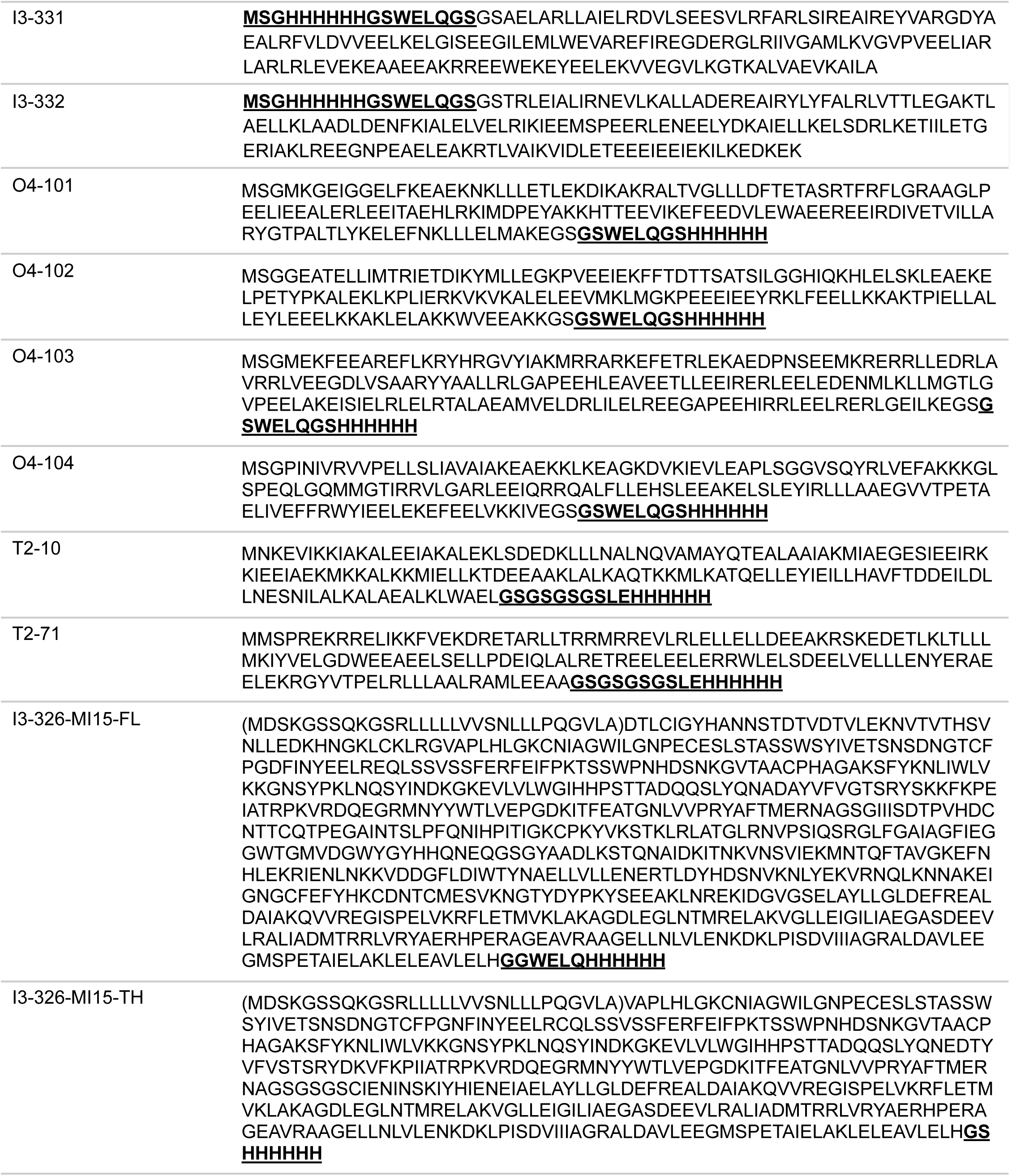
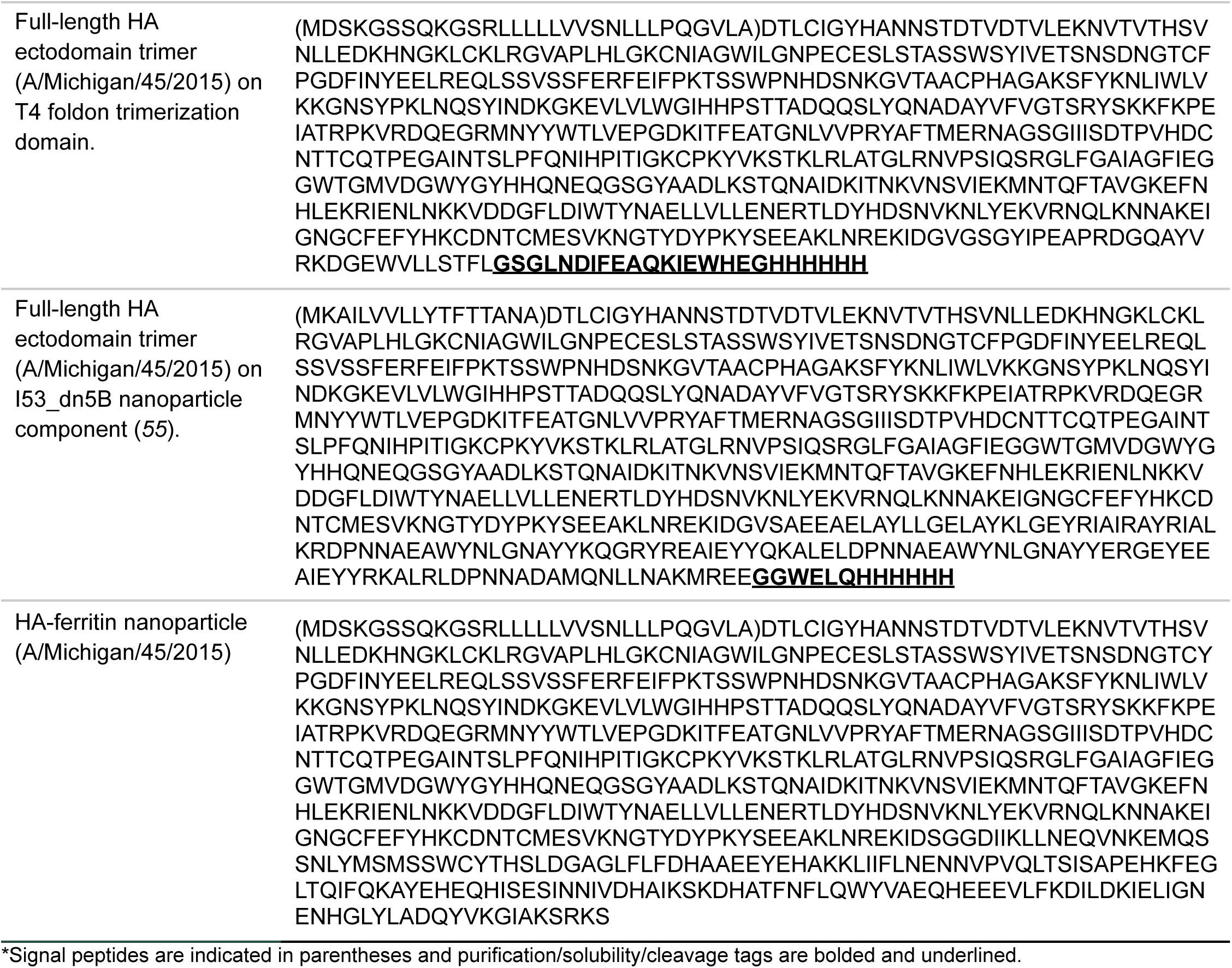
Amino Acid Sequences.

**Table S2.** CryoEM Data Collection, Processing, and Modeling Statistics (On-Target)

|  | **I3-304**<br>**PDB: 9ZQI**<br>**EMDB: 74566** | **O4-102**<br>**PDB: 9ZOJ**<br>**EMDB: 74498** | **O4-104**<br>**PDB: 9ZPM**<br>**EMDB: 74530** | **I3-326**<br>**PDB: 9ZSD**<br>**EMDB: 74706** |
| --- | --- | --- | --- | --- |
| **Data Collection** |  |  |  |  |
| Microscope | Titan Krios | Titan Krios | Titan Krios | Titan Krios |
| Voltage (kV) | 300 | 300 | 300 | 300 |
| Detector | Gatan K3 | Gatan K3 | Gatan K3 | Gatan K3 |
| Energy Filter | Gatan BioQuantum Gif | Gatan BioQuantum Gif | Gatan BioQuantum Gif | Gatan BioQuantum Gif |
| Recording mode | Counting | Counting | Counting | SuperResolution |
| Magnification | 105,000 X | 105,000 X | 105,000 X | 81,000 X |
| Movie micrograph pixel size (Å) | 0.83 | 0.83 | 0.83 | 0.5305 |
| Dose rate (e<sup>-</sup>/Å<sup>2</sup>/s) | 10.6 | 9.26 | 9.20 | 16.15 |
| Frames per movie micrograph | 99 | 99 | 99 | 75 |
| Frame exposure time (s) | 0.0505 | 0.0505 | 0.0505 | 0.0413 |
| Movie micrograph exposure time (s) | 5.009 | 5.009 | 5.009 | 3.096 |
| Total dose (e<sup>-</sup>/Å<sup>2</sup>) | 52.9 | 46.29 | 46.08 | 50 |
| Under focus range (µm) | 0.8 - 1.8 | 0.8 - 1.8 | 0.8 - 1.8 | 0.8 - 1.8 |
| No. of movie micrographs | 6,302 | 6,375 | 11,590 | 4,091 |
| **Map Processing** |  |  |  |  |
| Symmetry Applied | I | O | O | I |
| Extraction Box Size (pix) | 500 | 600 | 500 | 1400 |
| Initial particle images (no.) | 1,321,177 | 728,534 | 2,201,438 | 935,578 |
| Final particle images (no.) | 350,549 | 320,100 | 402,797 | 38,017 |
| Map resolution (Å) | 2.40 | 2.14 | 2.14 | 3.10 |
| FCS threshold | 0.143 | 0.143 | 0.143 | 0.143 |
| Map resolution range (Å) | 2.3 - 3.7 | 1.8 - 2.7 | 1.8 - 2.3 | 2.5-2.9 |
| **Refinement** |  |  |  |  |
| Initial model used | Design Model | Design Model | Design Model | Design Model |
| Map resolution (Å) | 2.40 | 2.14 | 2.14 | 3.10 |
| FCS threshold | 0.143 | 0.143 | 0.143 | 0.143 |
| Model resolution range (Å) | 2.35 - 2.50 | 1.8 - 2.4 | 1.8 - 2.3 | 2.5-2.9 |
| Map sharpening B factor | -115.1 | -76.9 | -83.0 | -127 |
| Model composition |  |  |  |  |
| Non-hydrogen atoms | 66,300 | 28,968 | 30,870 | 85,680 |
| Protein Residues | 9,000 | 3,600 | 3,600 | 11,100 |
| Ligands | Na | Na | Na | Na |
| Water | 2,161 | 2,556 | 2,022 | Na |
| B factors (Å) |  |  |  |  |
| Protein | 49.27 | 26.48 | 32.02 | 88.40 |
| Ligands | Na | Na | Na | Na |
| R.M.S. deviations |  |  |  |  |
| Bond lengths (Å) | 0.002 | 0.011 | 0.010 | 0.003 |
| Bond angles (°) | 0.445 | 1.798 | 1.265 | 0.561 |
| Validation |  |  |  |  |
| MolProbity score | 1.00 | 1.29 | 0.81 | 1.37 |
| Clashscore | 2.25 | 5.37 | 0.66 | 2 |
| Rotamer Outliers (%) | 0.75 | 0.00 | 1.32 | 3.43 |
| Ramachandran plot |  |  |  |  |
| Favored (%) | 99.07 | 99.44 | 99.27 | 99.33 |
| Allowed (%) | 0.60 | 0.56 | 0.73 | 0.13 |
| Disallowed (%) | 0.34 | 0.00 | 0.00 | 0.55 |

**Table S3.** CryoEM Data Collection, Processing, and Modeling Statistics (Additional I3-304 State)

| Off-Target-I3-304<br>PDB: 9ZOL<br>EMDB: 74499 |  |  |
| --- | --- | --- |
| Data Collection |  |  |
| Microscope | Titan Krios |  |
| Voltage (kV) | 300 |  |
| Detector | Gatan K3 |  |
| Energy Filter | Gatan BioQuantum Gif |  |
| Recording mode | Counting |  |
| Magnification | 105,000 X |  |
| Movie micrograph pixel size (Å) | 0.83 |  |
| Dose rate (e<sup>-</sup>/Å<sup>2</sup>/s) | 10.6 |  |
| Frames per movie micrograph | 99 |  |
| Frame exposure time (s) | 0.0505 |  |
| Movie micrograph exposure time (s) | 5.009 |  |
| Total dose (e<sup>-</sup>/Å<sup>2</sup>) | 52.9 |  |
| Under focus range (μm) | 0.8 - 1.8 |  |
| No. of movie micrographs | 6,302 |  |
| Map Processing |  |  |
| Symmetry Applied | D5 | C1 |
| Extraction Box Size (pix) | 500 |  |
| Initial particle images (no.) | 8,592 |  |
| Final particle images (no.) | 6,952 |  |
| Map resolution (Å) | 3.99 | 7.33 |
| FCS threshold | 0.143 |  |
| Map resolution range (Å) | 6.70 - 4.32 | 8.66 - 5.02 |
| Refinement |  |  |
| Initial model used | Design Model |  |
| Map resolution (Å) | 3.99 |  |
| FCS threshold | 0.143 |  |
| Model resolution range (Å) | 4.85 - 3.75 |  |
| Map sharpening B factor | 78.2 | 503.0 |
| Model composition |  |  |
| Non-hydrogen atoms | 33,720 |  |
| Protein Residues | 5,890 |  |
| Ligands | Na |  |
| *B* factors (Å) |  |  |
| Protein | 142.30 |  |
| Ligands | Na |  |
| R.M.S. deviations |  |  |
| Bond lengths (Å) | 0.003 |  |
| Bond angles (°) | 0.481 |  |
| Validation |  |  |
| MolProbity score | 1.20 |  |
| Clashscore | 4.17 |  |
| Rotamer Outliers (%) | 0.83 |  |
| Ramachandran plot |  |  |
| Favored (%) | 98.62 |  |
| Allowed (%) | 1.38 |  |
| Disallowed (%) | 0.00 |  |

**Table S4.** Crystallographic data collection and refinement statistics.

| **T2-71 (PDB: 9ZV3)** |  |
| --- | --- |
| Resolution range | 43.49 - 3.24 (3.42 - 3.24) |
| Space group | *R 3 2 :H* |
| Unit cell | 104.29, 104.29, 472.87; 90, 90, 120 |
| Unique reflections | 16281 (2302) |
| Multiplicity | 5.7 (5.9) |
| Completeness (%) | 100.00 (100.00) |
| Mean I/sigma(I) | 7.1 (1.1) |
| R-merge | 0.137 (1.420) |
| R-pim | 0.069 (0.709) |
| CC<sub>1/2</sub> | 0.996 (0.457) |
| Reflections used in refinement | 16274 (1314) |
| R-work | 0.1910 (0.3262) |
| R-free | 0.2356 (0.3728) |
| Number of non-hydrogen atoms |  |
| macromolecules | 5092 |
| solvent | 16 |
| protein residues | 600 |
| RMS(bonds) | 0.003 |
| RMS(angles) | 0.700 |
| Ramachandran favored (%) | 98.48 |
| Ramachandran allowed (%) | 1.52 |
| Ramachandran outliers (%) | 0.00 |
| Average B-factor |  |
| macromolecules | 107 |
| solvent | 68 |

## Notes

### Summary of Updates

Acknowledgements section updated; added remaining PDB IDs and EMDB IDs; supplementary table S2 was completed

